# Fatty acid binding protein-8 (FABP8/PMP2) reveals molecular heterogeneity of myelin sheaths in the human CNS

**DOI:** 10.64898/2026.06.09.730898

**Authors:** Vasiliki-Ilya Gargareta, Nikolaos Mougios, Elisabeth Teresa Hahn, Sophie B. Siems, Sarah J. Crisp, Balazs Varga, Ragnhildur Thora Karadottir, Joerg M. Buescher, Ramona B. Jung, Vidya Ramesh, Bhuvaneish T. Selvaraj, Siddharthan Chandran, Lida Zoupi, Jenea M. Bin, Maria A. Eichel-Vogel, David A. Lyons, Eneritz Agirre, Ting Sun, Gonçalo Castelo-Branco, Marina Uecker, Lars van Werven, Sabrina Zechel, Ulrike Fuchs, André Fischer, Wiebke Möbius, Ruth Stassart, Andrea J. Lopez, Petri Kursula, Olaf Jahn, Christine Stadelmann, Klaus-Armin Nave, Felipe Opazo, Hauke B. Werner

**Affiliations:** Department of Neurogenetics, Max Planck Institute for Multidisciplinary Sciences, Göttingen, Germany; Institute of Neuro- and Sensory Physiology, University Medical Center, Göttingen, Germany; Center for Biostructural Imaging of Neurodegeneration (BIN), University Medical Center, Göttingen, Germany; Department of Clinical Neurosciences, University of Cambridge, UK; Biosciences Institute, Newcastle University, Newcastle upon Tyne, UK; Cambridge Stem Cell Institute & Department of Veterinary Medicine, University of Cambridge, United Kingdom; Metabolomics Facility, Max Planck Institute of Immunobiology and Epigenetics, Freiburg, Germany; Centre for Clinical Brain Sciences, University of Edinburgh, Edinburgh, UK; UK Dementia Research Institute, University of Edinburgh, Edinburgh, UK; Institute for Neuroscience and Cardiovascular Research, University of Edinburgh, UK; Simons Initiative for the Developing Brain, University of Edinburgh, UK; MS Society Edinburgh Centre for MS Research, University of Edinburgh, UK; Laboratory of Molecular Neurobiology, Department of Medical Biochemistry and Biophysics, Science for Life Laboratory, Karolinska Institutet, Stockholm, Sweden; Neuroproteomics Group, Department of Molecular Neurobiology, Max Planck Institute for Multidisciplinary Sciences, Göttingen, Germany; Department of Neuropathology, University Medical Center, Göttingen, Germany; Department for Systems Medicine and Epigenetics, German Center for Neurodegenerative Diseases (DZNE), Göttingen, Germany; Electron Microscopy-City Campus, Max Planck Institute for Multidisciplinary Sciences, Göttingen, Germany; Paul Flechsig Institute, Center for Neuropathology and Brain Research, University of Leipzig, Leipzig, Germany; Department of Biomedicine, University of Bergen, Norway; Faculty of Biochemistry and Molecular Medicine, University of Oulu, Finland; Translational Neuroproteomics Group, Department of Psychiatry and Psychotherapy, University Medical Center, Göttingen, Germany; Department of Neuropathology, Charité, University Medical School, Berlin, Germany; Neurochemistry Group, Max Planck Institute for Multidisciplinary Sciences, Göttingen, Germany; Faculty for Biology and Psychology, University of Göttingen, Germany

**Keywords:** Oligodendrocyte, myelination, human CNS, white matter, myelin heterogeneity, myelin evolution, fatty acid binding protein 8 (FABP8/PMP2), ‘humanized’ mice, cholesterol, lipidomics

## Abstract

Myelin is classically viewed as a uniform axon-insulating membrane, yet its molecular composition may differ between species and even within one species. Fatty acid binding protein-8 (FABP8/PMP2) was previously identified in CNS myelin of humans but not mice. Here we show that FABP8/PMP2 is a defining feature of CNS myelin in humans and old-world-monkeys, but absent from CNS myelin in other mammals, indicating evolutionary neofunctionalization of this lipid-binding protein in the primate lineage. In the human CNS, FABP8/PMP2 marks a subset of myelin sheaths that preferentially ensheath large-diameter axons, revealing sheath-to-sheath molecular heterogeneity correlated with axonal geometry. Chromatin is accessible at the *PMP2/Pmp2* gene locus in oligodendrocytes of humans but not mice. Human oligodendrocytes intrinsically express FABP8/PMP2 when transplanted into mouse brains, demonstrating species-specific competence independent of environmental cues. ‘Humanized’ transgenic mice expressing FABP8/PMP2 in oligodendrocytes form morphologically normal but developmentally transiently thicker myelin sheaths, and show elevated cholesterol content in purified myelin. Because FABP8/PMP2 binds cholesterol, we propose that its emergence in primate CNS myelin contributes to the cholesterol enrichment of human myelin. Thus, CNS myelin protein composition is evolvable and modular, with relevance for myelin lipids and morphology, and previously unrecognized complexity in neuron–glia co-adaptation.

**Main points:**

- Fatty acid binding protein 8 (FABP8/PMP2) is present in CNS myelin of humans and old-world monkeys
- PMP2 defines sheath-to-sheath heterogeneity in the human CNS
- PMP2-immunopositive myelin ensheaths large-diameter axons
- Human oligodendrocytes intrinsically express PMP2 upon transplantation into mice
- ‘Humanized’ PMP2-transgenic mice show thicker myelin and altered myelin lipid composition

## Introduction

Oligodendrocytes enable normal motor, sensory and cognitive abilities of vertebrates via supporting neuronal axons in the central nervous system (CNS). Oligodendrocyte-neuron interactions include metabolic and antioxidative support, exosome transfer, and maintaining ion homeostasis upon axonal activity (Krämer-Albers and Werner, 2023; Stadelmann et al., 2019). Saltatory impulse propagation is facilitated by the ensheathment of axonal segments with myelin (Bunge et al., 1962; Hartline and Colman, 2007), a specialized multilayered extension of the oligodendroglial plasma membrane. The structure and functions of myelin require a particular molecular composition with specialized myelin proteins (Gargareta et al., 2022; Jahn et al., 2020) and lipids (Poitelon et al., 2020; Schmitt et al., 2015). It is commonly assumed that the unusually high lipid content of myelin (70-75%) is required for its property as an electrical insulator (O’Brien and Sampson, 1965). The functional relevance of healthy myelin sheaths is illustrated by the functional decline observed in human patients with myelin-related diseases including leukodystrophies and multiple sclerosis and respective animal models (Nowacki et al., 2022; Stadelmann et al., 2019; Wolf et al., 2020).

Myelin research is frequently performed in mice as a model for humans notwithstanding that species-dependent differences exist, including life span, body and brain size, and the lengths of axons. Given that the rodent and primate clades evolutionarily diverged about 85-90 million years ago (Murphy et al., 2001), we consider it relevant to know which aspects of myelin biology are conserved and which are specific to mice or humans. For example, it was previously recognized that myelin basic protein (MBP), which is essential for the formation of compact myelin in the CNS of mice (Readhead et al., 1987), displays one dominant isoform of 18.5 kDa in human but three main isoforms (14.0, 17.0, 18.5 kDa) in mouse CNS myelin (Ishii et al., 2009; Waehneldt and Malotka, 1980) due to species-dependent alternative splicing (Campagnoni, 1988), although the functional relevance has remained speculative.

More recently, it has become possible to systematically assess oligodendroglial transcriptome profiles in mice and humans, including in myelin-related disorders (Falcão et al., 2018; Pandey et al., 2022; Seeker et al., 2023; Zhou et al., 2020). The mRNA abundance profiles of healthy oligodendrocytes generally correlate well between both species; however, some clusters and distinct genes display species-dependent divergence (Gargareta et al., 2022; Jäkel et al., 2019). When the protein composition of myelin biochemically purified from brains of C57Bl/6N mice and subcortical normal appearing white matter (NAWM) of human individuals was compared using quantitative mass spectrometry, the relative abundance of structural myelin proteins (proteolipid protein (PLP), myelin basic protein (MBP), septins) was highly similar (Gargareta et al., 2022), suggesting evolutionary pressure and conservation. However, multiple other proteins were predominantly or exclusively identified in human or mouse myelin indicating heterogeneity of myelin protein composition across mammalian species. For example, tetraspanin-2 (TSPAN2) is a genuine CNS myelin protein in mice and rats (Birling et al., 1999; De Monasterio-Schrader et al., 2013; Jahn et al., 2020; Terada et al., 2002), but not present in human CNS myelin (Gargareta et al., 2022). Comparative analysis of myelin composition thus implies that mouse biology cannot be directly extrapolated to humans with respect to the function of TSPAN2 in myelin.

On the other hand, the presence of proteins in human but not mouse myelin (Gargareta et al., 2022) implies the existence of facets of myelin biology particularly relevant to humans. To investigate these further, we focused here on fatty acid binding protein-8 (FABP8), which by mass spectrometry was identified in human but not mouse CNS myelin (Gargareta et al., 2022; Jahn et al., 2020). Also termed peripheral myelin protein 2 (PMP2 or P2), FABP8 is a known constituent of PNS myelin synthesized by Schwann cells (Poitelon et al., 2020; Uusitalo et al., 2021) across reptile, bird, and mammalian species (Waehneldt et al., 1986). FABP8/PMP2 is a basic protein of 15 kDa that resides in the major dense line of compact PNS myelin (Poitelon et al., 2020). Mutations affecting the *PMP2* gene cause dominant demyelinating Charcot-Marie-Tooth disease (CMT type 1) (Gonzaga-Jauregui et al., 2015; Hong et al., 2016; Motley et al., 2016; Punetha et al., 2018). FABP8/PMP2 was previously found to bind cholesterol (Fang et al., 2024; Majava et al., 2010; Ruskamo et al., 2020; Sedzik and Jastrzebski, 2011). Apart from its cholesterol-binding properties, FABP8 may also associate with retinoic acid (Sedzik and Jastrzebski, 2011; Uyemura et al., 1984) or phosphatidylinositol (4,5)-bisphosphate (Abe et al., 2021), at least *in vitro*. FABP8/PMP2 can also affect lipid membrane morphology at least *in vitro*, including by stacking lipid bilayers into myelin-like multilayers (Krokengen et al., 2025; Ruskamo et al., 2020; Schöffmann et al., 2026; Uusitalo et al., 2021), in similarity to MBP.

This study aimed to characterize the expression and functional relevance of FABP8/PMP2 in CNS myelin. We found that FABP8/PMP2 is present in CNS myelin of several species of old-world monkeys including humans, but not of a variety of other mammalian species. In the human CNS, the presence of FABP8/PMP2 in myelin coincides with larger axonal diameters. Thus, human CNS myelin displays sheath-to-sheath heterogeneity. Finally, we established a transgenic mouse line in which human FABP8/PMP2 is expressed in oligodendrocytes and incorporated into CNS myelin; we propose that the consequences for the morphology of the axon/myelin-unit and myelin lipid composition observed in these mice reflect the functional relevance of FABP8/PMP2 in human myelin.

## Results

### Fatty acid binding protein-8 (FABP8/PMP2) is present in CNS myelin of old-world monkeys and humans

In a prior comparison of the CNS myelin proteome between humans and mice (Gargareta et al., 2022), FABP8/PMP2 was identified as CNS myelin protein specifically in humans. Here we first tested if presence of FABP8/PMP2 in CNS myelin extends beyond humans. To this end we used an established protocol (Erwig et al., 2019) to biochemically purify myelin from brain samples of various mammalian species, i.e. raccoon, pig, mouse, rat, marmoset, macaque, baboon, and human. To visualize the evolutionary relationships between these species, we plotted a phylogenetic tree using *Interactive Tree of Life* (<u>itol.embl.de</u>) (Letunic and Bork, 2024) (**Figure 1A**). The plot reflects that humans are more closely related to other primate species (marmoset, macaque, baboon) compared to rodents (mouse, rat), pig, and raccoon. By immunoblotting we detected FABP8/PMP2 in myelin purified from human, macaque, and baboon brains, but not in the other assessed species (**Figure 1B**). This suggests that FABP8/PMP2 is a CNS myelin protein in the primate clade Catarrhini (old-world monkeys and humans), but not in other mammalian clades including Platyrrhini (new-world monkeys).

**Figure 1.**
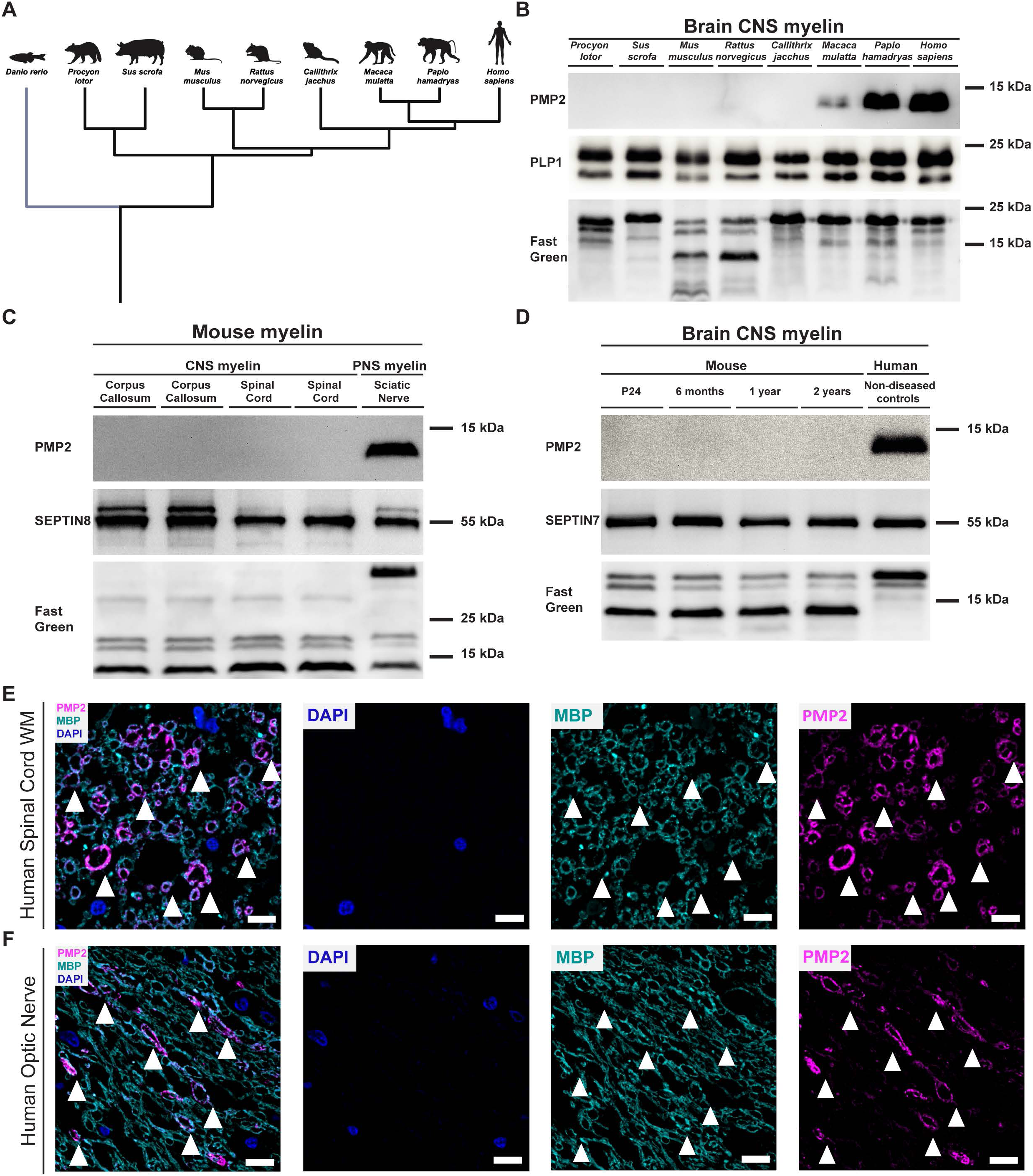
Fatty acid binding protein 8 (FABP8/PMP2) is present in CNS myelin of humans and old-world monkeys. **A,B** Phylogenetic tree **(A)** schematically depicting evolutionary relations between species analyzed by immunoblot **(B)** of myelin purified from the brains of *procyon lotor* (raccoon), *sus scrofa* (pig), *mus musculus* (mouse), *rattus norvegicus* (rat), *callithrix jacchus* (marmoset), *macaca mulatta* (rhesus macaque), *papio hamadryas* (baboon) and *homo sapiens* (human). **A** Phylogenetic tree generated with iTOL.embl.de. Danio rerio (zebrafish) is included in the phylogenetic tree as an outgroup. Sketched with Biorender. **B** Immunoblot shows presence of FABP8/PMP2 in myelin of multiple species of old-world monkeys and humans. The myelin marker PLP serves as positive control and Fast Green staining serves as loading control. Immunoblot shows one sample per species. **C** Immunoblot analysis detects FABP8/PMP2 in myelin purified from the peripheral nervous system (sciatic nerve) at P21 but not in myelin purified from the CNS (corpus callosum, spinal cord) of mice at P18. SEPTIN8 serves as positive control and Fast Green staining serve as loading control. Immunoblot shows n=2 biological replicates for corpus callosum and spinal cord and n=1 biological replicate for sciatic nerve. **D** Immunoblotting detects FABP8/PMP2 in myelin purified from human NAWM in the brain but not in myelin purified from brains of mice at P24, 6 mo, 12 mo, and 24 mo. SEPTIN7 serves as positive control and Fast Green staining serves as loading control. Immunoblot shows n=1 biological replicate per condition. **E-F** Immunofluorescence labelling and confocal microscopy of FABP8/PMP2 (magenta) and MBP (myelin marker; cyan) on cross-sectioned spinal cord **(E)** and optic nerve **(F)** of human individuals. Note that PMP2 immunolabelling is confined to a subset of MBP-immunopositive myelin sheaths (arrowheads point at double-immunopositive myelin rings). Nuclear staining by DAPI in blue. Micrographs show one individual, representative of n=3 **(E,F)** individuals. Scale bars 10 μm.

So far, the statement that FABP8/PMP2 is absent from CNS myelin in mice has largely been based on mass spectrometric and immunoblot analysis of myelin purified from young adult mouse brains. We thus used immunoblotting to test if FABP8/PMP2 is detected in myelin of mouse brains during development or ageing, or in another CNS region, i.e. the spinal cord. However, no FABP8/PMP2-signal was detected in myelin purified from spinal cords (**Figure 1C**) or from the brains of mice at P24, 6 months, 1 year, and 2 years (**Figure 1D**). Myelin fractions purified from the peripheral sciatic nerve of a mouse (**Figure 1C**), or from the brain of a human individual (**Figure 1D**), were used as respective positive controls. The latter experiment also confirms that the antibody is suited for detecting FABP8/PMP2 by immunoblot in both species. These data support the finding that FABP8/PMP2 is a myelin protein in the human CNS but not in the mouse CNS.

### Sheath-to-sheath heterogeneity of FABP8/PMP2 immunolabeling in the human CNS

To further characterize the expression of FABP8/PMP2 in the human CNS, we performed immunofluorescence labeling on cross-sectioned spinal cords (**Figure 1E**) and optic nerves (**Figure 1F**) of non-diseased individuals. By confocal microscopy, FABP8/PMP2 immunolabeling was visualized in both CNS regions as ring-like structures (white arrowheads in **Figure 1E-F**), suggesting its localization in myelin. Notably, FABP8/PMP2 immunolabeling was confined to a subset of myelin sheaths as identified by co-immunolabelling the myelin marker myelin basic protein (MBP; green in **Figure 1E-F**). For comparison, immunofluorescence labeling on cross-sectioned corpora callosa **(Supplemental Figure S1A)** or spinal cords (**Supplemental Figure S1B**) of mice and confocal microscopy did not detect FABP8/PMP2 immunolabeling in the mouse CNS, concurrent with immunoblotting (**Figure 1C-D**). Conversely, when using cross-sectioned peripheral sciatic nerves of mice as a positive control, FABP8/PMP2 immunolabeling was readily detected in a subset of MBP-immunopositive PNS myelin sheaths (**Supplemental Figure S1C**) in agreement with prior reports (Trapp et al., 1979; Yim et al., 2022), also confirming that the antibody is suited for detecting FABP8/PMP2 by immunolabeling of nervous tissue sections in both species. Together, these data show that FABP8/PMP2 is a constituent of human CNS myelin, in which its expression displays sheath-to-sheath heterogeneity.

### FABP8/PMP2 expression in human CNS myelin correlates with larger axonal diameters

Given that human myelin sheaths display sheath-to-sheath heterogeneity of FABP8/PMP2-expression (**Figure 1E-F**), we asked if this correlates with properties of the axon/myelin-unit. To this end, we used stimulated emission depletion (STED) microscopy to analyze immunofluorescence labelling of FABP8/PMP2 and the myelin marker PLP on cross-sectioned optic nerves of eight non-diseased human individuals (**Figure 2A**). In agreement with confocal microscopy (**Figure 1F**), this visualized myelin rings double-immunopositive for FABP8/PMP2 and PLP, as well as rings single-immunopositive for PLP. We then determined the inner boundary of PLP-immunopositivity as a proxy of axonal diameter and the outer boundary of PLP-immunopositivity as a proxy for the diameter of the axon/myelin-unit and calculated their ratio (g-ratio) (**Figure 2B,C**). However, the g-ratio did not differ between axon/myelin-units immunopositive or immunonegative for FABP8/PMP2. Interestingly, the scatter plot of the g-ratio versus axonal diameters (**Figure 2C**) suggested a difference in the diameters of axons myelinated with sheaths that were immunopositive or immunonegative for FABP8/PMP2. Indeed, when assessing median axonal diameters and the relative frequency distribution (**Figure 2D,E**), axons with FABP8/PMP2 immunopositive sheaths displayed larger diameters compared to those with FABP8/PMP2 immunonegative sheaths in optic nerves. Similar results were obtained upon immunofluorescence labelling of FABP8/PMP2 and PLP using STED microscopy of cross-sectioned spinal cords from nine non-diseased human individuals (**Figure 2F-J**) in which FABP8/PMP2 immunopositive myelin sheaths also did not differ from FABP8/PMP2 immunonegative sheaths with respect to g-ratio (**Figure 2G,H**), but by ensheathing axons with larger calibers (**Figure 2I,J**). Axonal diameters with FABP8/PMP2-immunopositive or FABP8/PMP2-immunonegative sheaths did not differ grossly across individuals (**Supplemental Figure S2A,B**), suggesting that the correlation between larger axonal diameters and FABP8/PMP2-positivity of their myelin sheaths is not owing to a subset of individual samples. When using linear quantile mixed models (Bin et al., 2025; Geraci, 2014) to assess the diameters of axons, the diameters of axons with FABP8/PMP2-immunopositive sheaths differed significantly from those with FABP8/PMP2-immunonegative sheaths (**Supplemental Figure S2A,B**). By comparison of means across quartile groups, the difference was significant in all quartiles (**Supplemental Figure S2C,D**), suggesting that the correlation between larger axonal diameters and FABP8/PMP2-positivity of their myelin sheaths is not owing to a particular axon diameter range. Together, these data indicate that in humans, at least in the optic nerve and spinal cord, larger-caliber axons are preferentially associated with myelin sheaths expressing FABP8/PMP2.

**Figure 2.**
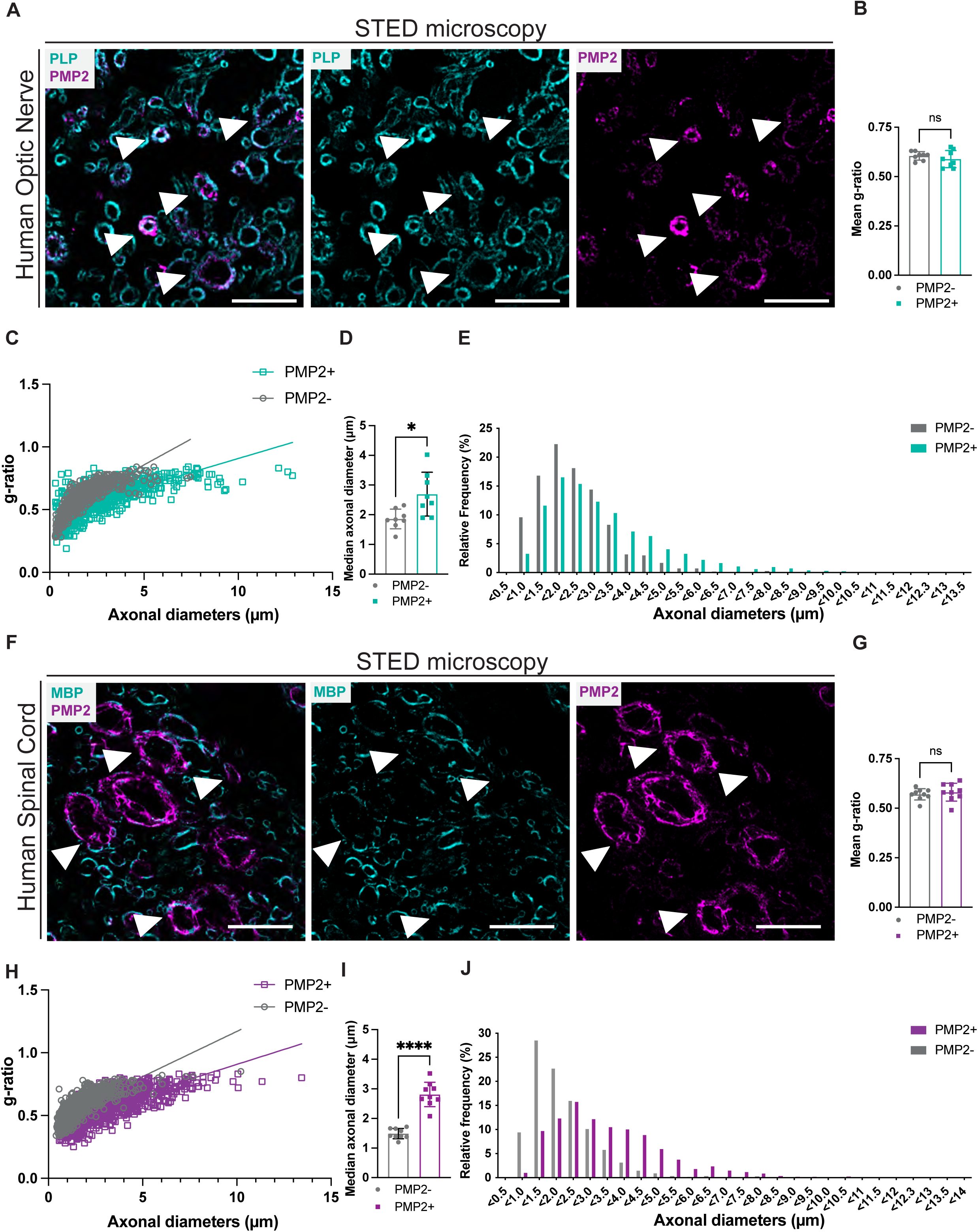
PMP2 is present in myelin ensheathing larger-diameter axons in humans. **A** Immunofluorescence labelling and superresolution microscopy (STED) showing PMP2 (magenta) and PLP (cyan; myelin marker) on cross-sectioned optic nerves of non-diseased human individuals. Arrowheads point at myelin rings double-immunopositive for PLP and PMP2. Scale bars 5 μm. **B,C** g-ratio analysis of micrographs as in **A** shows no difference in the ratio between axonal diameters and the diameters of the respective axon/myelin-unit comparing PMP2-immunopositive and PMP2-immunonegative myelin sheaths, suggesting appropriate myelin sheath thickness. Datapoints represent human individuals in **B** and axon/myelin-units in **C**. Optic nerves from n=8 non-diseased human individuals with 35-160 axons per sample. Bar graph in **B** shows mean ±SD; n.s. p>0.05 by Two-tailed Student’s t-test. **D,E** PMP2 immunopositivity-dependent quantification of micrographs as in **A** reveals larger diameters of axons with PMP2-immunopositive sheaths in human optic nerves. **D** Bar graph shows mean ±SD; datapoints represent human individuals. *p=0.0113 by Two-tailed Student’s t-test. **E** Frequency distribution with 0.5 μm bin width; n=8 individuals with 873 PMP2-immunopositive and 532 PMP2-immunonegative sheaths. **F** Immunofluorescence labelling and superresolution microscopy (STED) showing PMP2 (magenta) and MBP (cyan; myelin marker) on cross-sectioned spinal cords of non-diseased human individuals. Arrowheads point at myelin rings double-immunopositive for MBP and PMP2. Scale bar 5 μm. **G,H** g-ratio analysis of micrographs as in **F** shows similar ratios between axonal diameters and the diameters of the respective axon/myelin-unit comparing PMP2-immunopositive and PMP2-immunonegative myelin sheaths, suggesting appropriate myelin sheath thickness. Datapoints represent human individuals in **G** and axon/myelin-units in **H**. n=9 spinal cords from non-diseased human individuals with 37-171 axons per sample. Bar graph in **G** shows mean ±SD; n.s. p>0.05 by Two-tailed Student’s t-test. **I,J** PMP2 immunopositivity-dependent quantification of micrographs as in **F** reveals larger diameters of axons with PMP2-immunopositive sheaths in spinal cords. **I** Bar graph shows mean ±SD; datapoints represent human individuals; **** p<0.0001 by Two-tailed Student’s t-test. **J** Frequency distribution with 0.5 μm bin width; n=9 individuals with 1276 PMP2-immunopositive and 717 PMP2-immunonegative sheaths.

### The *PMP2/Pmp2* gene locus is accessible in human but not mouse oligodendrocytes

We next used dot plots to visualize the abundance of *PMP2/Pmp2* mRNA in myelin-forming oligodendrocytes (MOL in **Figure 3A**) and oligodendrocyte precursor cells (OPC in **Supplemental Figure S3A**) of brains and cervical spinal cords (CSC) according to previously published single cell multiomics datasets, i.e. single-nucleus RNA sequencing (snRNA-seq) and single-cell Assay for Transposase-Accessible Chromatin using sequencing (scATAC-seq) of humans (Kabbe et al., 2026) and mice (Bravo González-Blas et al., 2023; Zheng et al., 2025). Indeed, *PMP2*/*Pmp2* mRNA is present in a substantial percentage of both MOL and OPC in humans but not in mice **(Figure 3A**, **Supplemental Figure S3A)**. For comparison, transcripts encoding the mouse-specific (Gargareta et al., 2022) myelin protein TSPAN2 are present in a high percentage of both MOL and OPC in mice but not humans, transcripts encoding the marker of the oligodendrocyte lineage SOX10 (SRY-related HMG-box 10) (Sock and Wegner, 2021) are present in MOL and OPC of both species, transcripts encoding the MOL marker MYRF (myelin regulatory factor) (Emery and Wood, 2024) are more prominently expressed in MOL than in OPC, and transcripts encoding the OPC marker PDGFRA (platelet derived growth factor receptor α) (Emery and Wood, 2024) are largely confined to OPC **(Figure 3A**, **Supplemental Figure S3).** Thus, both the single-nucleus (sn)RNA-seq based data **(Figure 3A**, **Supplemental Figure S3)** and a prior single-cell (sc)RNA seq-based comparison (Gargareta et al., 2022) show that presence of FABP8/PMP2 in human but not mouse myelin correlates well with the expression of its transcripts in human but not mouse MOL.

**Figure 3.**
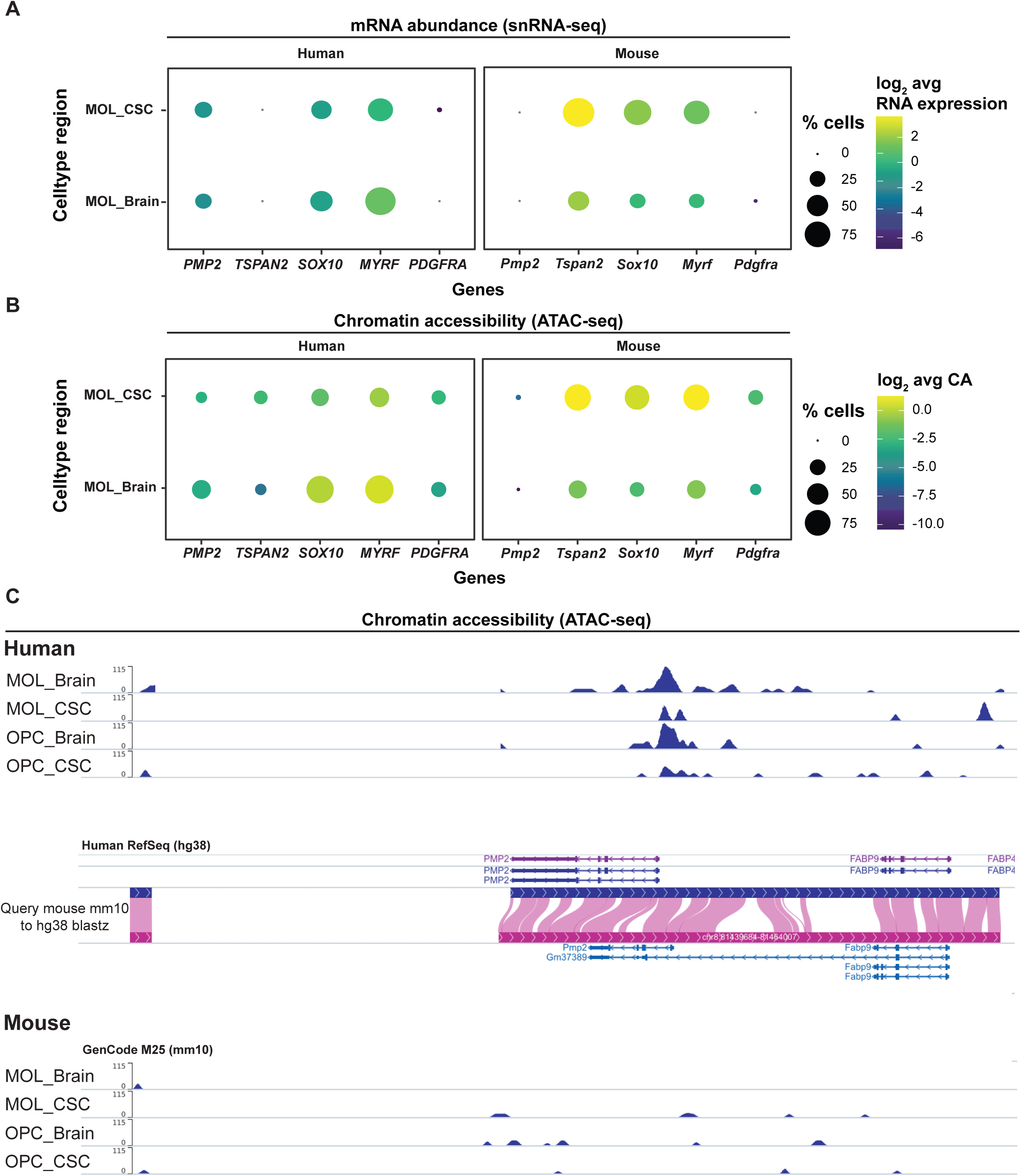
Cross-species snRNA-seq and scATAC-seq analysis of *PMP2/Pmp2* in oligodendrocytes. **A** Dot plot showing the percentage of myelinating oligodendrocytes (MOL) expressing *PMP2/Pmp2* and the indicated marker mRNAs, and their log_2_-transformed average mRNA abundance, in the cervical spinal cord (CSC) and brains. Data are taken from previously established human (left) and mouse (right) multiome single-nucleus RNA sequencing (snRNA-seq) datasets (Bravo González-Blas et al., 2023; Kabbe et al., 2026; Zheng et al., 2025). Note that *PMP2/Pmp2* is expressed in a considerable percentage of human but not mouse MOL. For oligodendrocyte precursor cells (OPC) see **supplemental Figure S3A**. **B** Dot plot showing the percentage of myelinating oligodendrocytes (MOL) in which the gene locus of *PMP2/Pmp2* and the indicated marker genes is accessible, and their log_2_-transformed average chromatin accessibility, in the cervical spinal cord (CSC) and brains. Data are taken from previously established human (left) and mouse (right) single-cell Assay for Transposase-Accessible Chromatin using sequencing (scATAC-seq) datasets (Bravo González-Blas et al., 2023; Kabbe et al., 2026; Zheng et al., 2025). Note that the *PMP2/Pmp2* locus is accessible in a considerable percentage of human but not mouse MOL. For oligodendrocyte precursor cells (OPC) see **supplemental Figure S3B**. **C** Plot showing accessible chromatin regions of the *PMP2*/*Pmp2* locus in MOL and OPC in brains and CSC of humans and mice. Peaks indicate relative accessibility. Note that the human *PMP2/Pmp2* locus is accessible in MOL and OPC of both brains and CSC at and around the transcription start site, indicating an active promoter region. In comparison, peaks are smaller or absent in mice.

We hypothesized that the species-dependent expression of *PMP2/Pmp2* is due to transcriptional differences between human and mouse oligodendrocytes. Activity of the *PMP2/Pmp2* gene is regulated by the transcription factor SOX10 (Graf et al., 2019; Williams et al., 2023). Indeed, sequence motifs indicating binding sites for the transcription factor SOX10 and its cofactor NFATC2 are present in the promoter region of both the human and the mouse *PMP2/Pmp2* gene within 1000 bp upstream and 100 bp downstream from the transcription initiation site according to analysis using the R package *motifmatchr* (v1.28.0) (Alicia Schep [Aut, 2017). To assess if *PMP2/Pmp2* mRNA expression in the oligodendrocyte lineage correlates with chromatin accessibility, we visualized accessibility of the gene locus in MOL (**Figure 3B**) and OPC (**Supplemental Figure S3B)** of brains and CSC of humans and mice according to previously published scATAC-seq datasets (Bravo González-Blas et al., 2023; Kabbe et al., 2026; Zheng et al., 2025). Indeed, by dot plot visualization the *PMP2*/*Pmp2* locus is accessible in a substantial percentage of both MOL and OPC in humans but not in mice **(Figure 3B**, **Supplemental Figure S3B)**. For comparison, the locus encoding the mouse-specific myelin protein TSPAN2 is accessible in a higher percentage of MOL and OPC in brains and CSC of mice compared to humans. It was interesting to note that chromatin accessibility at the loci encoding the markers SOX10, MYRF, and PDGFRA in MOL and OPC was not entirely similar when comparing human and mouse MOL and OPC **(Figure 3B**, **Supplemental Figure S3B)**. Notwithstanding that other variables may also play a role, this suggests that the gene regulatory network in the oligodendrocyte lineage displays some degree of species-dependent variability.

We also plotted open chromatin regions of the *PMP2*/*Pmp2* locus in MOL and OPC in brains and in the CSC of both humans and mice (**Figure 3C**). The positions of the peaks indicate a high degree of open chromatin regions in both cell types of both CNS regions of humans mainly upstream of the transcription initiation site, indicating an active promoter region. According to the Genome Reference Consortium Human Build 38 (GRCh38), the transcription initiation site of the human *PMP2* gene on chromosome 8 is located at position chr8:81447439 while the respective scATAC peak in oligodendrocytes spans chr8:81446971-81448578, suggesting that the human *PMP2* gene promoter as inferred from the ATAC signal extends 1138 bases upstream of the PMP2 transcription initiation site. Peaks were smaller or absent in OPC and MOL in brains and CSC of mice (**Figure 3C**). Together, it is plausible that the presence of PMP2 in human but not mouse CNS myelin is due to species-specific transcript expression in oligodendrocytes, which depends on species-dependent chromatin accessibility of the locus.

### Human oligodendrocytes express FABP8/PMP2 upon transplantation into mouse brains

To assess if expression of FABP8/PMP2 is an intrinsic characteristic of human oligodendrocytes or requires induction by the environment in the human brain, we first used a chimeric model in which dissociated glial spheres composed of human embryonic stem cell (hESC)-derived glial cells, including oligodendrocyte progenitor cells (OPCs), were transplanted into neonatal immunodeficient mice that lack compact CNS myelin (*Mbp^shi/shi^;Rag1^−/−^* mice) as previously reported (Ramesh et al., 2025). Upon transplantation, human OPCs were allowed to differentiate for 12 weeks *in vivo* before immunofluorescence labelling analysis of coronally sectioned corpora callosa. Given that *Mbp^shi/shi^*mice lack MBP-expression (Readhead et al., 1987), any MBP-immunolabelling signal detected derives from transplanted human oligodendrocytes. By immunofluorescence labelling we detected a number of cells double-immunopositive for MBP and FABP8/PMP2 (**Figure 4A**). This shows that human oligodendrocytes express FABP8/PMP2 upon transplantation into neonatal mouse brains, suggesting that FABP8/PMP2-expression is an intrinsic characteristic of human oligodendrocytes independent of their environment in the human brain.

**Figure 4.**
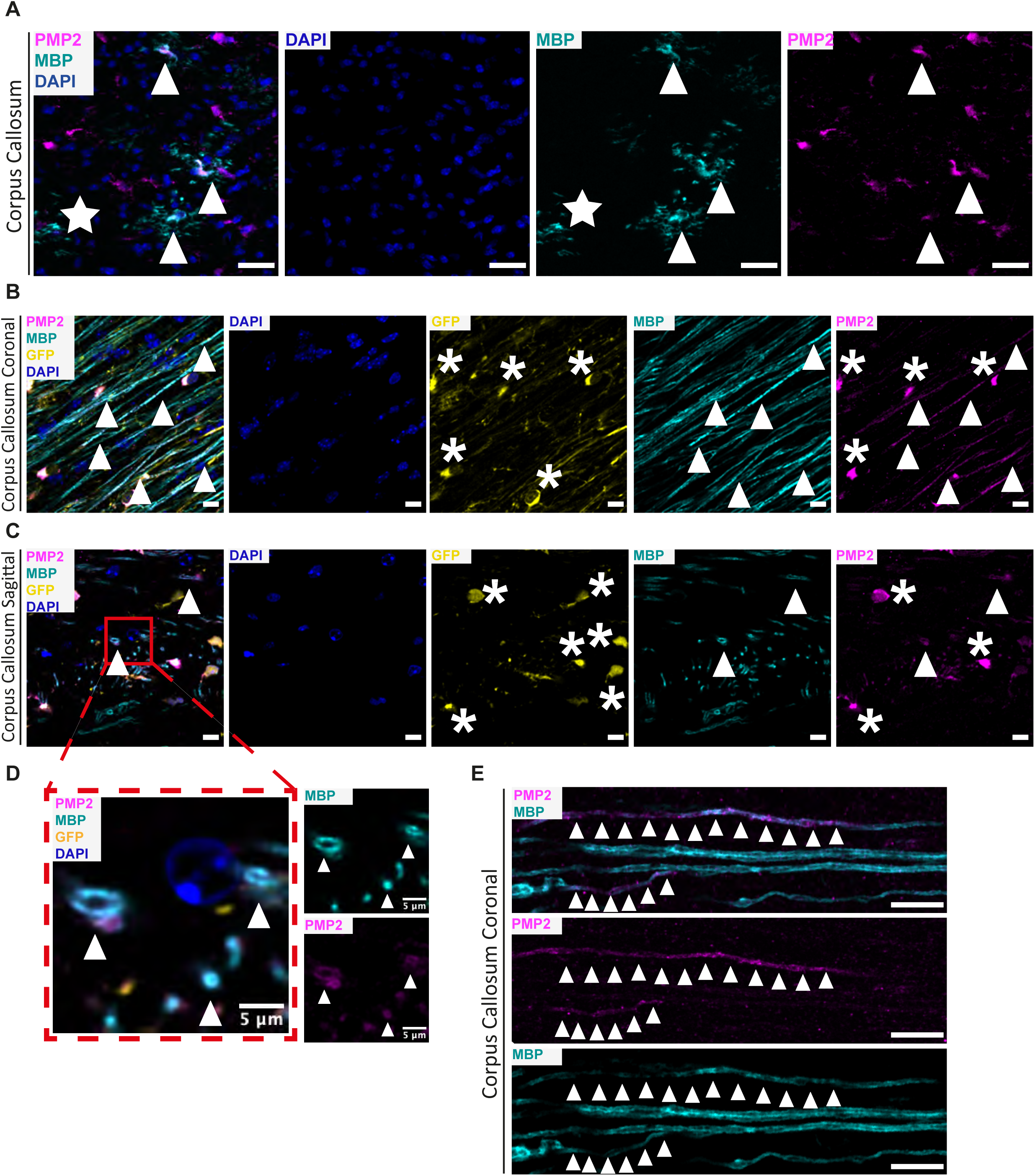
PMP2 is present in human-derived oligodendrocytes and myelin sheaths upon transplantation into mice. **A** Immunofluorescence labelling and confocal microscopy showing PMP2 (magenta) and MBP (myelin marker; cyan) labelling and DAPI (blue) on coronally sectioned corpora callosa of *Mbp^shi/shi^;Rag1^−/−^*mice at age at 12 weeks, which were injected with human embryonic stem cell (hESC) derived-derived oligodendrocytes at neonatal age P0-P1 as previously reported (Ramesh et al., 2025). Note the presence of both PMP2^+^/MBP^+^ double-immunopositive (white arrowheads) and PMP2^-^/MBP^+^ single-immunopositive (white star) oligodendrocytes. Micrograph representative of n=3 biological replicates. Scale bar 20 μm. **B-E** Immunofluorescence labelling and confocal microscopy showing PMP2 (magenta) and MBP (cyan) labelling and DAPI (blue) on one coronal and one sagittal section each of corpora callosa of *Myrf^flox/flox^;FoxG1^Cre+/WT^;Rag2^-/-^*mice at age 6 months, which were transplanted at age P0 with GFP-labelled human iPSC-derived oligodendrocytes (white asterisks highlighting yellow cell bodies). Micrograph representative of n=3 biological replicates. Note the presence of GFP^+^/PMP2^+^ double-positive oligodendrocyte cell bodies (white asterisks) and PMP2^+^/MBP^+^ double-immunopositive myelin sheaths (white arrowheads). **B** Coronal section of the corpus callosum providing a view of myelin sheaths along the axon. Scale bar 10 μm. **C-D** Sagittal section of the corpus callosum providing a view of cross-sectioned myelin sheaths. **C** Scale bar 10 μm. **D** Magnification of red box in **C**; Scale bar 5 μm. **E** Magnified view of coronally sectioned corpus callosum shows both PMP2^+^/MBP^+^ double-immunopositive (white arrowheads) and PMP2^-^/MBP^+^ single-immunopositive myelin sheaths, reflecting sheath-to-sheath heterogeneity of PMP2-expression. Scale bar 10 μm.

In this model, transplanted human oligodendrocytes did not generate fully extended, mature myelin sheaths. We thus employed a second *in vivo* chimeric model, in which OPCs derived from human induced pluripotent stem cells (iPSC) expressing green fluorescent protein (GFP) were neonatally transplanted into immunodeficient mice lacking myelination in the forebrain including the corpus callosum (*Myrf^flox/flox^;FoxG1^Cre+/WT^;Rag2^-/-^*mice). Given that GFP is not expressed in mouse cells, MYRF is essential for myelination, and FoxG1-Cre mediates recombination of floxed alleles in the forebrain (Collins et al., 2025; Emery et al., 2009; Kawaguchi et al., 2016), all GFP-positive cells and most myelin sheaths in the forebrain of this model derive from transplanted iPSC-derived human oligodendrocytes. We then performed immunofluorescence labeling of FABP8/PMP2 and MBP on coronally (**Figure 4B,E**) and sagitally sectioned (**Figure 4C,D**) corpora callosa. By confocal microscopy, we identified cells double-positive for GFP and FABP8/PMP2 (highlighted with asterisks in **Figure 2B,C**), as well as longitudinal (**Figure 4B**) and ring-like (**Figure 4C**) structures double-immunopositive for MBP and FABP8/PMP2 (arrowheads in **Figure 4B,C**). Higher magnifications highlight the presence of ring-like structures upon sagittal sectioning (arrowheads in **Figure 4D**) and longitudinal structures upon coronal sectioning (arrowheads in **Figure 4E**) of corpora callosa double-immunopositive for MBP and FABP8/PMP2. We interpret these findings as reflecting the presence of mature human oligodendrocytes that extend FABP8/PMP2 immunopositive myelin sheaths upon transplantation into mouse brains. We also noted the presence of structures single-immunopositive for MBP but immunonegative for FABP8/PMP2, which may reflect sheath-to-sheath heterogeneity of its expression (**Figure 1D-F**, **Figure 2**). It will be an interesting topic of future research to assess if myelin sheaths of human oligodendrocytes transplanted into mouse brains adopt a relationship between PMP2-immunopositivity and axonal diameters. Together, these data imply that the expression of FABP8/PMP2 and its incorporation into myelin are intrinsic features of human oligodendrocytes independent of their environment in the human brain, and that sheath-to-sheath heterogeneity is maintained upon transplantation into mice.

### A ‘humanized’ mouse line expressing FABP8/PMP2 in oligodendrocytes

To assess the consequences of *de novo* expression of FABP8/PMP2 in CNS myelin, we generated a transgenic mouse line in which the human FABP8/PMP2 coding sequence is specifically expressed in myelinating glia. Expression is driven by the CAG promoter (Jun-ichi et al., 1989); cell type-specificity is mediated by the presence of a stop sequence (3* polyA signal) flanked by loxP sites that are recombined in myelinating cells upon interbreeding with mice expressing Cre-recombinase under control of the *Cnp* promotor (*Cnp^Cre/WT^*) (Lappe-Siefke et al., 2003) (Scheme in **Figure 5A**). Indeed, *tg^CAG-loxP-Stop-loxP-Pmp2^*;*Cnp^Cre/WT^* mice were achieved and in the following are termed PMP2OE (for *Pmp2^overexpressor^*). To assess if FABP8/PMP2 is present in myelin of PMP2OE mice, we biochemically purified myelin from their brains. By immunoblotting (**Figure 5B**), FABP8/PMP2 was readily detected in PMP2OE myelin while no signal was detected in myelin purified from the brains of *tg^CAG-loxP-Stop-loxP-Pmp2^* mice without the presence of Cre, which served as control (Ctrl) mice.

**Figure 5.**
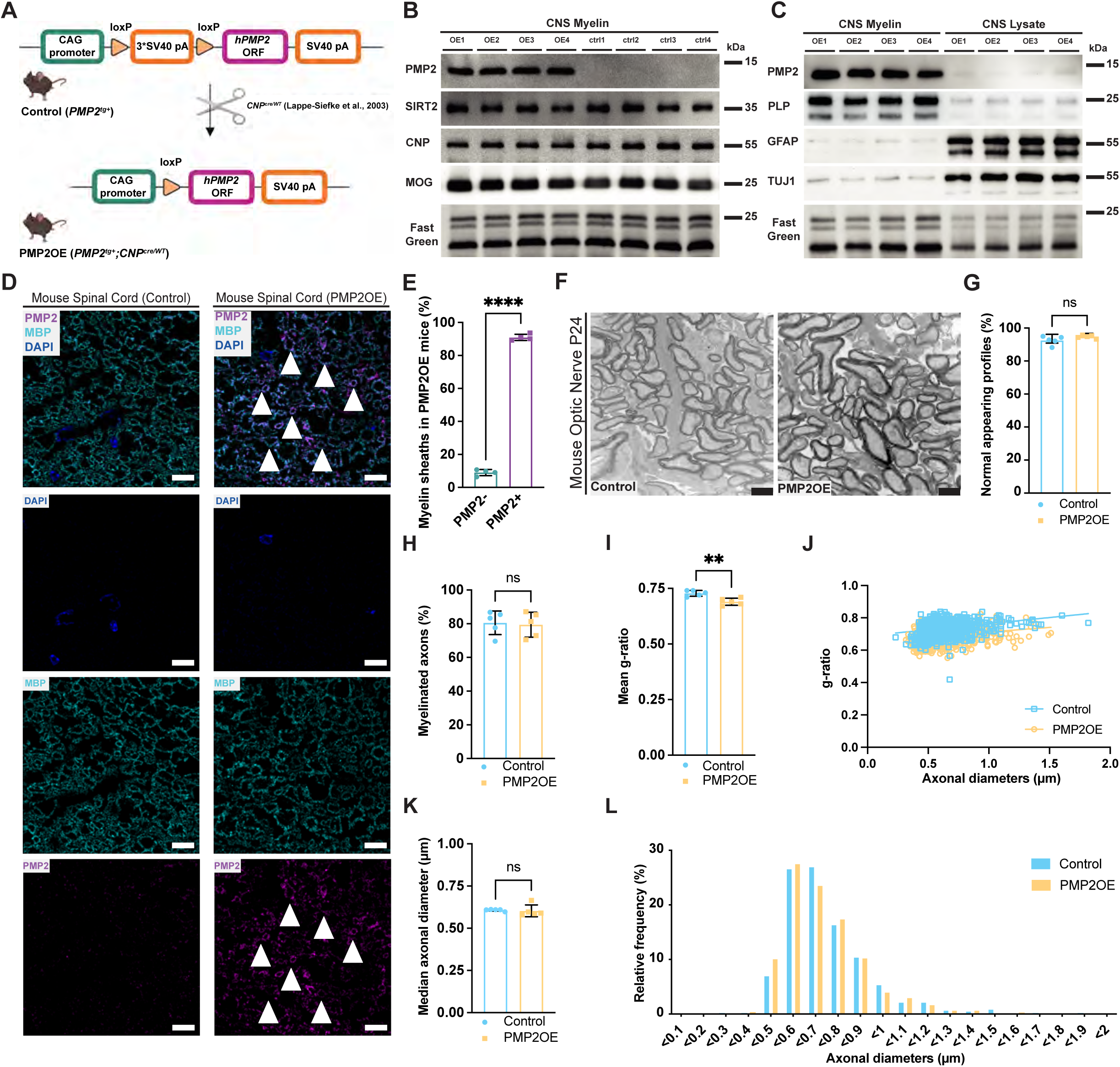
‘Humanized’ mice expressing *hPMP2* in mouse myelinating glia (PMP2OE) **A** Scheme of transgene construct for PMP2 ‘humanized’ mice. Construct comprises the CAG promoter, a stop sequence (3* SV40 polyA signals) flanked by loxP sites, the coding sequence of *hPMP2*, and another polyA signal. Transgene-positive mice were interbred with mice expressing Cre-recombinase under control of the *Cnp* promoter (*Cnp^Cre/WT^*) (Lappe-Siefke et al., 2003) gaining *hPmp2^tg+^;Cnp^Cre/WT^* mice, also termed PMP2OE. Sketched with Biorender. **Β** Immunoblot analysis showing presence of PMP2 in myelin purified from the brains of PMP2OE (OE1-OE4) but not myelin of control (ctrl 1-ctrl 4) mice. Myelin markers SIRT2, CNP, and MOG were detected as controls. Fast Green staining serves as loading control. Blots show n=4 biological replicates per genotype. **C** Immunoblot analysis showing enrichment of PMP2 in myelin purified from the brains of PMP2OE mice (OE1-OE4) compared to respective brain lysate. Markers for myelin (PLP), astrocytes (GFAP), and axons (TUJ1) were detected to assess the enrichment of myelin proteins and the concomitant depletion of astrocytic and axonal proteins. Fast Green staining serves as loading control. Blots show n=4 biological replicates. **D** Immunofluorescence labelling and confocal microscopy of PMP2 (magenta) and MBP (cyan) on cross-sectioned spinal cord of PMP2OE and control mice at 9 months. DAPI in blue. Note that PMP2-immunopositive rings (arrowheads) are readily detectable in PMP2OE but not in control mice. Images representative of n=4 mice per genotype. Scale bar 10 μm. **E** Quantitative assessment of PMP2 immunopositive (PMP2+) and immunonegative (PMP2-) myelin sheaths in PMP2OE mice (n=4) on images as in **D**. Bar graph; mean ±SD; datapoints represent individual mice; ****p<0.0001 by Two-tailed Student’s t-test. Note that most myelin sheaths in spinal cords of PMP2OE mice are PMP2-immunopositive. **F-L** Electron micrographs of cross-sectioned optic nerves dissected from control and PMP2OE mice at P24 **(F)** and genotype-dependent quantification **(G-L)**. **F** Images representative of n=5 mice per genotype. Scale bar 1 μm. **G** Quantification of non-pathological axon/myelin-units on images as in **F** reveals no significant difference between control and PMP2OE mice at P24 (n=5). Mean ±SD; datapoints represent individual mice; n.s. by Two-tailed Student’s t-test. **H** Quantification of the percentage of axons that are myelinated on images as in **F** reveals no significant difference between control and PMP2OE mice at P24 (n=5). Mean ±SD; datapoints represent individual mice; n.s. by Two-tailed Student’s t-test. **I,J** g-ratio analysis of normal appearing axon-myelin profiles in optic nerves reveals a decreased g-ratio suggesting thicker myelin sheaths in PMP2OE compared to Ctrl mice at P24 (n=5). **I** Bar graph gives mean ±SD; ** p=0.0032 by Two-tailed Student’s t-test. **J** Scatter plot displays g-ratio and respective axonal diameters; datapoints show 85-172 axon/myelin-units per mouse. **K,L** Genotype-dependent quantification reveals similar axonal diameters in optic nerves of PMP2OE and Ctrl mice at P24 (n=5). **K** Bar graph gives mean ±SD; n.s. p>0.05 by Two-tailed Student’s t-test. **L** No frequency distribution shift in PMP2OE mice compared to controls. Data presented as frequency distribution with 0.1 μm bin width; 556 axons in n = 5 control and 700 axons in n = 5 PMP2OE mice.

Next, we used immunoblotting to compare the abundance of marker proteins in purified myelin and brain lysate of PMP2OE mice (**Figure 5C**). We found that FABP8/PMP2 was markedly enriched in CNS myelin of PMP2OE mice, similar to the myelin marker PLP. On the other hand, the astrocyte marker GFAP and the axonal marker TUJ1 (also termed TUBB3) were strongly reduced in myelin compared to the brain lysate of PMP2OE mice (**Figure 5C**), reflecting the degree of purity of the myelin fraction. We also used immunofluorescence labelling and confocal microscopy to examine the expression of FABP8/PMP2. Myelin sheaths were recognized by co-immunolabeling MBP. FABP8/PMP2 was readily detectable in the spinal cords of PMP2OE mice (**Figure 5D**), while no signal was detected in control mice. When quantifying myelin sheaths that were double-immunopositive for MBP and FABP8/PMP2 in PMP2OE spinal cords, we found that over 85% of MBP-immunopositive sheaths were also immunopositive for FABP8/PMP2 (**Figure 5E**). Only few MBP-immunopositive myelin sheaths in PMP2OE spinal cords were PMP2-immunonegative (**Figure 5E**). Together, this suggests that the CAG promoter is active in oligodendrocytes, loxP site recombination takes place, and human FABP8/PMP2 is efficiently incorporated into CNS myelin of PMP2OE mice. We thus considered PMP2OE mice a suitable model to assess the functional relevance of FABP8/PMP2 in CNS myelin.

To this end, we next used transmission electron microscopy to analyse the ultrastructure of axon/myelin-units in optic nerves dissected from juvenile PMP2OE and control mice at P24 (**Figure 5F**). By quantitative evaluation of electron micrographs, we found a similar number of normal-appearing (i.e., non-pathological) axon/myelin-units in PMP2OE and control mice (**Figure 5G**), indicating that human FABP8/PMP2 can be expressed in mouse myelin without causing pathology of axons or myelin. For example, pathological myelin outfoldings or whorls were not a feature. We also found a similar percentage of axons being myelinated (**Figure 5H**); however, the g-ratio was decreased in PMP2OE compared to control mice (**Figure 5I,J**). This suggests that expression of FABP8/PMP2 in PMPOE mice causes myelin sheaths that are thicker than appropriate for the respective axonal diameters at this age. When assessing the diameters of myelinated axons (**Figure 5K,L**), also using linear quantile mixed models (Bin et al., 2025; Geraci, 2014) (**Supplemental Figure S4A**) and comparison of means across quartile groups (**Supplemental Figure S4B)**, there was no genotype-dependent effect. To assess the ultrastructure of axon/myelin-units at a more advanced age, we turned to electron microscopy of optic nerves dissected from young adult PMP2OE and control mice at P45 (**Supplemental Figure S4C**). By quantitative evaluation of electron micrographs, we found no evidence of enhanced pathology of axon/myelin-units (**Supplemental Figure S4D**) and no change in the percentage of axons being myelinated (**Supplemental Figure S4E**) in PMP2OE mice at this age, in similarity to juvenile mice (**Figure 5G,H**). Moreover, when assessing the g-ratio of axon-myelin-units we found no difference between PMP2OE and control mice (**Supplemental Figure S4F,G**). This implies that the increased myelin sheath thickness in juvenile PMP2OE mice (**Figure 5I,J**) is a transient, developmental effect. When assessing the diameters of myelinated axons (**Supplemental Figure S4H-K**), we found similar axonal diameters in PMP2OE compared to control mice. Together, this suggests that oligodendroglial expression of PMP2 can increase CNS myelin sheath thickness transiently in young mice; however, we did not find evidence that expression of PMP2 in myelin causes increased axonal diameters.

### Proteome and lipidome analysis of CNS myelin in PMP2OE mice

We hypothesized that the introduction of a lipid-binding protein into CNS myelin of PMP2OE mice may affect its protein or lipid composition. To systematically compare the protein composition of CNS myelin between PMP2OE and control mice, we biochemically purified myelin from brains dissected at P45 and performed quantitative mass spectrometry using a previously described workflow that includes two data acquisition modes, i.e. MS^E^ for the correct quantification of exceptionally abundant myelin proteins (PLP, MBP, CNP) and UDMS^E^ for deep quantitative coverage of medium-to-low abundant myelin proteins (Gargareta et al., 2022; Jahn et al., 2020; Siems et al., 2020). As expected, FABP8/PMP2 was exclusively identified in CNS myelin of PMP2OE but not of control mice, with a sequence coverage of 43%, including several unique peptides specific to human FABP8/PMP2 (**Figure 6A; Supplemental Data Table S1**), in agreement with immunoblot (**Figure 5B**). All other known myelin proteins were identified and quantified in myelin of both genotypes; none of them displayed a considerable difference in abundance in myelin between the genotypes **(Figure 6A; Supplemental Figure S5A,B, Supplemental Data Table S1)**. For example, structural myelin proteins including PLP, MBP, and myelin septins displayed similar abundance in PMP2OE and control myelin. This indicates that incorporation of FABP8/PMP2 into CNS myelin of mice does not have significant secondary consequences for its protein composition.

**Figure 6.**
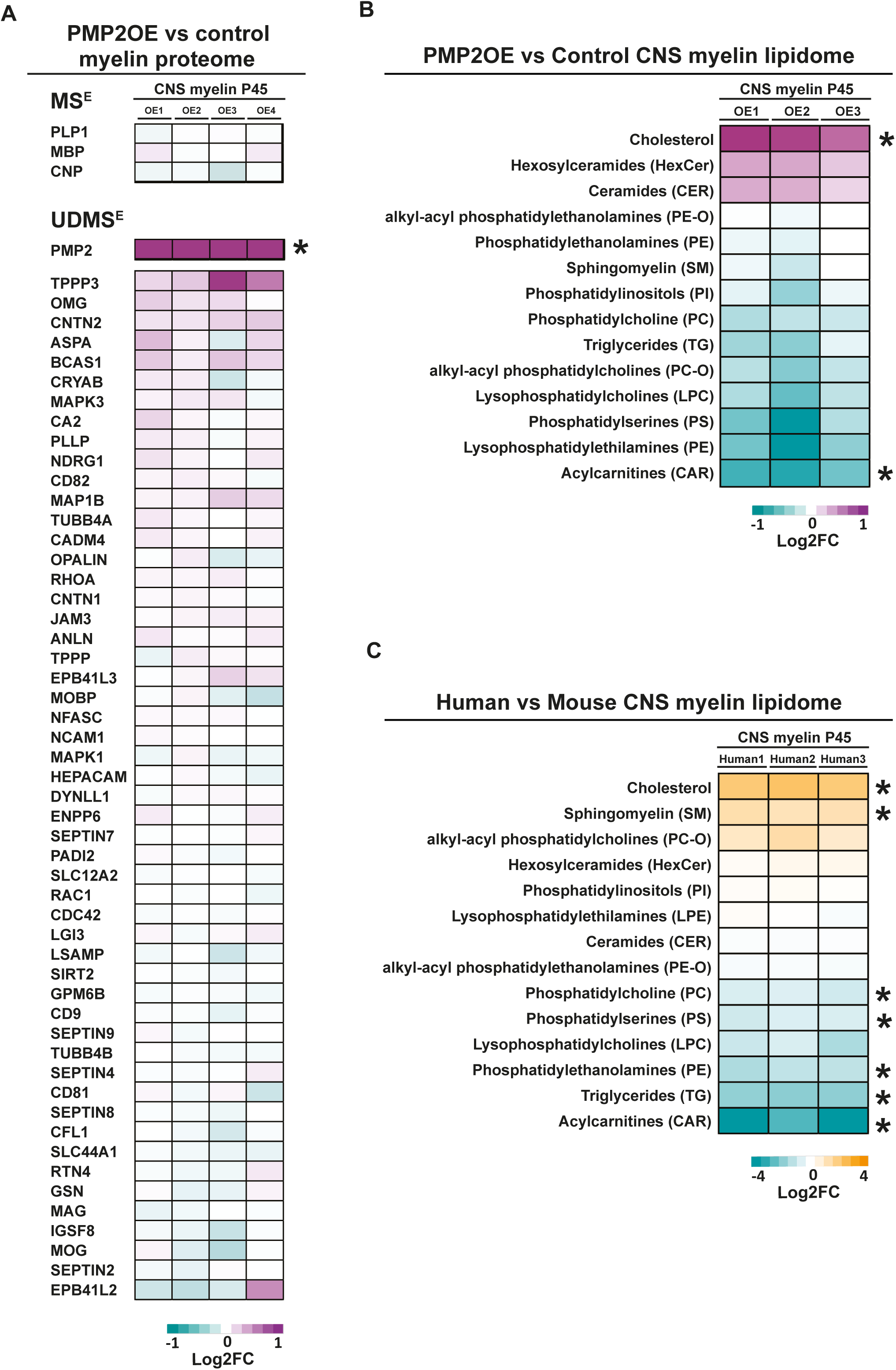
Myelin proteome and lipidome analysis of PMP2OE mice and comparison of myelin lipids between mouse and human brains. **A** Differential proteome analysis comparing the relative abundance of proteins in myelin purified from brains of PMP2OE and control mice at P45 (n=4 per genotype). Heatmaps show mass spectrometric quantification of known myelin proteins. Each horizontal line displays the fold-change (FC) of a myelin protein in 4 biological replicates of PMP2OE myelin (OE1-OE4) as the average of two technical replicates each compared to the mean abundance in control myelin, plotted on a log_2_-color scale. Magenta represents higher abundance in PMP2OE, cyan represents higher abundance in control myelin. Exceptionally abundant myelin proteins (PLP. MBP, CNP) and other myelin proteins are assessed using measurement modes MS^E^ and UDMS^E^ respectively. Note that, except for PMP2, no known myelin protein displays genotype-dependent difference in abundance. For volcano plots of dataset see **Supplemental Figure S6**. For entire dataset see **Supplemental Data Table S1**. **B** Differential mass spectrometric analysis of lipids in myelin purified from the brains of PMP2OE (magenta) and control (cyan) mice at P45 (n=3) using positive ionization mode. Heat map shows the relative abundance of the indicated lipid classes in myelin. Each horizontal line displays the fold-change (Log2FC) of a lipid class in 3 biological replicates of PMP2OE myelin (OE1-OE3) compared to the mean abundance in control myelin, plotted on a log_2_-color scale. Magenta represents higher abundance in PMP2OE, cyan represents higher abundance in control myelin. Statistical analysis via multiple unpaired t-test followed by Sidak-Bonferroni correction. Stars indicate significant difference; Cholesterol *p=0.0018, CAR *p=0.0027. Note the higher abundance of cholesterol in PMP2OE compared to control myelin. For volcano plot showing individual lipid species see **Supplemental Figure S6C**; for dataset see **Supplemental Data Table S2**. **C** Differential mass spectrometric analysis of lipids in myelin purified from normal-appearing subcortical white matter of non-diseased humans (n=3) and C57Bl/6N mouse brains (n=3) using positive ionization mode. Heat map shows the relative abundance of the indicated lipid classes in myelin. Each horizontal line displays the fold-change (Log2FC) of a lipid class in 3 biological replicates of human myelin (Human1-Human3) compared to the mean abundance in mouse myelin, plotted on a log_2_-color scale. Orange represents higher abundance in human myelin, cyan represents higher abundance in mouse myelin. Statistical analysis via multiple unpaired t-test followed by Sidak-Bonferroni correction. Stars indicate significant difference. Cholesterol *p=0.0001, SM *p=0.00002, PC *p=0.0003, PS *p=0.0014, PE *p=0.00002, TG *p=0.0005, CAR *p=0.0001. Note the higher abundance of cholesterol in human compared to mouse myelin. For entire dataset see **Supplemental Data Table S2.**

Considering that FABP8/PMP2 is a fatty acid binding protein we next hypothesized that its incorporation into CNS myelin may affect the lipid composition of the latter. We thus used mass spectrometry to compare the lipid composition of CNS myelin purified from PMP2OE and control mice at P45. As expected, lipid species were readily identified in myelin of both genotypes using both positive and negative ionization modes (**Supplemental Data Table S2**). Most lipid classes displayed similar abundance in myelin of PMP2OE and control mice (**Figure 6B, Supplemental Figure S5C**). The relative abundance of acetyl-carnitines (CAR) was moderately but significantly decreased in PMP2OE compared to control myelin. Most noticeably, however, the relative abundance of cholesterol was considerably increased in PMP2OE compared to control myelin (**Figure 6B, Supplemental Figure S5C**). Given that FABP8/PMP2 is a cholesterol-binding protein (Fang et al., 2024; Majava et al., 2010; Ruskamo et al., 2020; Sedzik and Jastrzebski, 2011), this suggests that FABP8/PMP2 either facilitates cotransport of cholesterol into – or retains it in – the myelin sheath, thus contributing to its lipid composition.

### Comparative analysis of lipids in CNS myelin of mice and humans

The finding that presence of FABP8/PMP2 affects the relative abundance of lipids in CNS myelin of mice motivated us to compare the lipid composition of CNS myelin between humans and mice. We note that the species-dependent differences between mice and humans in the transcriptional profile of oligodendrocytes and the proteome of CNS myelin (Gargareta et al., 2022; Jäkel et al., 2019; Seeker et al., 2023) that may affect its lipid composition, are not limited to FABP8/PMP2. Yet, we subjected myelin biochemically purified from the subcortical white matter of non-diseased human individuals (comprising FABP8/PMP2) and the brains of Ctrl mice (lacking FABP8/PMP2) to quantitative mass spectrometric analysis. Lipids were identified mode in human and mouse myelin using both positive and negative ionization mode **(Supplemental Data Table S2)**. A number of lipid classes was detected as more abundant in mouse compared to human myelin, including carnitines (CAR), phosphatidylethanolamine (PE), phosphatidylserine (PS), and phosphatidylcholine (PC) **(Figure 6C; Supplemental Figure S5D)**. On the other hand, multiple lipid species were more abundant in human compared to mouse myelin, including sphingomyelin (SM). Most prominently, cholesterol was identified as more abundant in human compared to mouse CNS myelin **(Figure 6C; Supplemental Figure 5D)**, in similarity to the comparison between PMP2OE and control CNS myelin (**Figure 6B, Supplemental Figure S5C**).

## Discussion

The existence of differences in the proteomes of CNS myelin between humans and mice implies that some aspects of myelin biology are specific to a subgroup of mammalian species. By mass spectrometric comparison of myelin protein composition (Gargareta et al., 2022), we previously identified FABP8/PMP2 as a constituent of human but not mouse CNS myelin. By immunoblot analysis we here find that FABP8/PMP2 is present in CNS myelin of multiple species of old-world monkeys including humans but not in several other mammalian species including a new-world monkey, notwithstanding that the number of assessed species was limited by availability. It is possible that FABP8/PMP2 was already present in CNS myelin of now-extinct species at the root of mammals, and, after evolutionary divergence from the clade including old-world monkeys and humans, was eliminated by selective constraints in other mammals. However, we consider it more plausible that FABP8/PMP2 was clade-specifically neofunctionalized as a CNS myelin protein at the root of the clade including old-world monkeys and humans by recruitment from more ancient functions, e.g. in Schwann cells and PNS myelin (Belin et al., 2019; Hong et al., 2024; Poitelon et al., 2020; Uusitalo et al., 2021; Zenker et al., 2014). This hypothesis is supported by our finding that FABP8/PMP2 can be transgenically expressed in mouse oligodendrocytes and is incorporated into their myelin sheaths without causing evident neuropathology, which argues against the existence of selective constraints. It is probable that evolutionary emergence of oligodendroglial *PMP2* expression in ancient old-world monkeys involved evolutionary changes in chromatin accessibility at the *PMP2* locus.

### Sheath-to-sheath heterogeneity of human CNS myelin

When immunolabelling human CNS tissue and detecting FABP8/PMP2 by STED microscopy, we unexpectedly found heterogeneity of its immunopositivity across myelin sheaths in multiple CNS regions. The analysis of previously published multiomics snRNA-seq and scATAC-seq datasets (Bravo González-Blas et al., 2023; Kabbe et al., 2026; Zheng et al., 2025) showed that the *PMP2* locus is accessible and *PMP2* mRNA is expressed in only a subset of oligodendrocytes. It is thus plausible that the observed sheath-to-sheath heterogeneity is owing to the existence of oligodendrocyte subtypes that can be distinguished by *PMP2*-expression. Interestingly, the presence of FABP8/PMP2 in human myelin sheaths correlates with larger axonal diameters. This suggests that either the presence of FABP8/PMP2 in myelin may induce large-diameter axons, or that diameter-dependent axonal signals affect myelination by oligodendrocytes including those that express FABP8/PMP2. We did not find evidence that incorporation of FABP8/PMP2 into CNS myelin of transgenic mice induces a shift towards larger diameters of myelinated axons, and, more generally, myelination of CNS axons *per se* does not affect their diameters in healthy developmental conditions (Bin et al., 2025). On the other hand, our finding that human oligodendrocytes transplanted into mouse brains can express FABP8/PMP2 and display sheath-to-sheath heterogeneity suggests that any relevant axonal signals are not only present in humans but also in mice. Yet, our data do not exclude the existence of a yet-unknown mechanism by which axonal subtypes differing in their surface signals locally affect the molecular composition of the myelin sheaths that myelinate them. To the best of our knowledge this is the first report of sheath-to-sheath heterogeneity of the expression of a myelin protein, at least in humans. Yet, our data imply the possibility that other CNS myelin proteins with sheath-to-sheath heterogeneity of their expression remain to be discovered.

Interestingly, PNS myelin synthesized by Schwann cells also displays molecular sheath-to-sheath heterogeneity. Indeed, neighboring peripheral sheaths differ with respect to FABP8/PMP2 expression in both mice and humans, and the presence of FABP8/PMP2 in PNS myelin correlates with larger axonal diameters (Trapp et al., 1979; Yim et al., 2022), thus reminding of central sheaths in humans. It is thus tempting to speculate that the mechanisms responsible for sheath-to-sheath heterogeneity in CNS and PNS may be similar. In the PNS, a subtype of Schwann cells expressing *Pmp2* transcripts myelinate larger-diameter axons, mainly motor axons (Yim et al., 2022). Interestingly, this Schwann cell population is required for the long-term maintenance of the axons they myelinate, as shown by their diphtheria-toxin-induced ablation (Kozlowski et al., 2025). Yet it remains unknown if the mechanisms by which these Schwann cells maintain the survival of peripheral axons involve FABP8/PMP2 itself (Kozlowski et al., 2025); it appears more likely that FABP8/PMP2 is a mere marker for the relevant Schwann cell population.

In the CNS, subtypes of oligodendrocytes have been identified according to mRNA expression profile clusters, which depend on CNS region, sex, developmental state, ageing (Marisca et al., 2020; Marques et al., 2016; Seeker et al., 2023), and disease conditions (Depp et al., 2023; Falcão et al., 2018; Jäkel et al., 2019). While some oligodendrocyte subtypes are characterized by the expression of genes relevant for axonal support, others are defined by diverging expression of myelination-related genes (Jäkel et al., 2019). Interestingly, a recent comparison of oligodendroglial mRNA profiles across multiple CNS regions in humans revealed differences in the frequency of occurrence of oligodendrocyte subtypes between the human brain and spinal cord (Seeker et al., 2023). One subcluster defined by elevated expression of *PMP2* mRNA was enriched in human spinal cord (Seeker et al., 2023), possibly reflecting that this CNS region is particularly rich in large-diameter axons. Together, the presence of FABP8/PMP2 in myelin likely correlates with subtypes of Schwann cells in the PNS and subtypes of oligodendrocytes in the human CNS respectively.

On the other hand, our work does not exclude the possibility that signals derived from central large-diameter axons may affect oligodendroglial expression of FABP8/PMP2. For example, axonal neuregulin (NRG1 type III) does not only affect proliferation of and myelination by Schwann cells via the PI3K/AKT and MAPK/ERK pathway (Brinkmann et al., 2008; Meyer and Birchmeier, 1995; Michailov, 2004; Taveggia et al., 2005), but also enhances transcription of the *Pmp2* gene with NRG1 type III-dependent consequences for myelination in the PNS (Belin et al., 2019; Della-Flora Nunes et al., 2025; Hong et al., 2024). In the CNS, lack of NRG1 impairs oligodendrocyte differentiation *in vitro* (Vartanian et al., 1999) while its overexpression stimulates myelination when combined with brain-derived neurotrophic factor (BDNF) (Brinkmann et al., 2008; Lundgaard et al., 2013). While neuregulin or other axonal factors may possibly affect oligodendroglial FABP8/PMP2 expression, our transplantation experiment indicates that the principal competence to express FABP8/PMP2 is a feature specifically of human (but not mouse) oligodendrocytes. Conversely, possible relevant axonal signal(s) (if any) are unlikely to be limited to humans. It will be an interesting topic of future investigation to test if any among the many known axon-to-oligodendrocyte signals (Adams et al., 2021; Ahrendsen and Macklin, 2013; Nave and Werner, 2014; Stadelmann et al., 2019) can stimulate oligodendroglial *PMP2* expression.

### A ‘humanized’ mouse model expressing FABP8/PMP2 in CNS myelin

Transgenic expression of human FABP8/PMP2 in oligodendrocytes of mice causes temporarily increased myelin thickness during developmental myelination. In the PNS, the thickness of myelin sheaths is unchanged in non-injured mice lacking FABP8/PMP2 expression (Hong et al., 2024; Uusitalo et al., 2021; Zenker et al., 2014). On the other hand, overexpression of FABP8/PMP2 in cultured Schwann cells increases the number and length of myelinated segments of dorsal root ganglia neurons *in vitro* (Della-Flora Nunes et al., 2025), and mice lacking FABP8/PMP2 display transiently thinner peripheral myelin sheaths during remyelination in the sciatic nerve crush injury paradigm *in vivo* (Zenker et al., 2014). Notwithstanding that effects of loss-of-function and gain-of-function can principally differ, as may those in Schwann cells and oligodendrocytes, these data imply that expression of FABP8/PMP2 has myelination-promoting effects in both CNS and PNS, possibly via its cholesterol-binding properties (Fang et al., 2024; Majava et al., 2010; Sedzik and Jastrzebski, 2011).

Compared to other biomembranes, myelin membranes contain an unusually high percentage (70-75%) of lipids with an approximate ratio 2:2:1:1 of cholesterol, phospholipids, galactolipids and plasmalogens (Norton and Poduslo, 1973). Considering that prospective myelin membranes expand by the coalescence of cholesterol-rich membrane microdomains in the secretory pathway of the myelinating cell (Chrast et al., 2011; Lee, 2001; Simons et al., 2000), it is not surprising that cholesterol availability affects myelin biogenesis. Indeed, mutant mice or zebrafish that lack an enzyme essential for cholesterol biosynthesis display impaired myelination (Mathews et al., 2014; Saher et al., 2005). It is thus plausible that the temporarily increased myelin thickness during development is explained by the expression of an additional myelin protein and, more speculatively, the increased availability of cholesterol in CNS myelin of mice upon expression of FABP8/PMP2. Indeed, similar to MBP, FABP8/PMP2 binds lipid membranes at least *in vitro* (Krokengen et al., 2025; Ruskamo et al., 2020; Schöffmann et al., 2026; Uusitalo et al., 2021), thereby affecting the morphology of myelin-like multilayers with an intermembrane distance of 1 nm larger than with MBP alone. It is of note, however, that FABP8/PMP2 is not the only cholesterol-binding myelin protein that contributes to myelin lipid composition. Indeed, the most abundant protein constituents of central and peripheral myelin, PLP and myelin protein zero (MPZ/P0), respectively, and probably multiple other myelin proteins bind cholesterol and may thus contribute to the cholesterol level in myelin by binding and cotransport (Ruskamo et al., 2022; Saher et al., 2009; Sedzik et al., 2013; Simons et al., 2000; Werner et al., 2013). The presence or absence of FABP8/PMP2 thus does not define, but may rather modulate, the cholesterol content of myelin beyond a basic level.

### Potential disease relevance of PMP2 in the CNS

*PMP2* is a known disease gene for the peripheral neuropathy Charcot-Marie-Tooth disease (CMT1) (Gonzaga-Jauregui et al., 2015; Hong et al., 2016; Motley et al., 2016; Punetha et al., 2018), which is not surprising considering that it has long been recognized as a myelin-related gene and myelin protein in the PNS of rodents (Brostoff et al., 1974; Trapp et al., 1979; Uusitalo et al., 2021; Waehneldt et al., 1986; Zenker et al., 2014). Our data imply that it is a relevant task for future research to assess if CMT1 patients with causative PMP2 mutations display CNS-based dysfunction. More recently, it was found that *PMP2* displays transcriptional changes in patients with Alzheimer’s Disease (Kelley et al., 2018), and that FABP8/PMP2 binds amyloid-β (Aβ) fibers *in vitro* and associates with Aβ plaques *in vivo* according to immunofluorescence labelling (Montero-Calle et al., 2023), notwithstanding that the precise involvement of FABP8/PMP2 in Alzheimer’s Disease remains to be further scrutinized. In view of our identification of FABP8/PMP2 as a myelin protein in the human CNS, and its prior identification in human (but not mouse) astrocytes (Kelley et al., 2018), we suggest FABP8/PMP2 as a plausible candidate to be considered in human myelin-related disorders of the CNS, which are now recognized to include Alzheimer’s Disease (Chen et al., 2021; Depp et al., 2023; Rajani et al., 2024; Sasmita et al., 2024).

### Limitations of the study

Our ‘humanized’ mice expressing FABP8/PMP2 in CNS myelin did not model sheath-to-sheath heterogeneity as observed in the human CNS, given the very high percentage of FABP8/PMP2-immunopositive sheaths. On the other hand, the latter enabled biochemical characterization of CNS myelin and thus the identification of a correlation between FABP8/PMP2 positivity and cholesterol levels. Both the presence of FABP8/PMP2 (Schöffmann et al., 2026) and varying cholesterol levels (Maxfield and Van Meer, 2010; Rosetti et al., 2008; Saher et al., 2009; Yurlova et al., 2011; Zhu et al., 2026) might alter the membrane properties of myelin. To dissect their relative contributions was beyond the scope of the present project but represents an interesting topic of future research. We also note that the experimental lack of FABP8/PMP2 from Schwann cells (Zenker et al., 2014) affects the lipid composition of peripheral myelin at least during early postnatal development, with the relative abundance of sphingomyelin and phosphatidylcholine decreased and that of glycosphingolipids increased. However, cholesterol was not indicated in that dataset (Zenker et al., 2014), making it impossible for us to judge if its abundance in PNS myelin correlates with the presence or absence of FABP8/PMP2.

## Conclusion

The protein composition of CNS myelin differs between mammalian groups at least with respect to the lipid-binding protein FABP8/PMP2, which is present in myelin sheaths in the CNS of old-world monkeys, including humans, but not other mammalian groups. Our work did not reveal evidence of selective constraints against expression of FABP8/PMP2 in mouse CNS myelin, suggesting that it was neofunctionalized at the root of the old-world monkey clade. The competence to express FABP8/PMP2 is an intrinsic feature of human oligodendrocytes, also when transplanted into mouse brains. Our expression analysis demonstrates that human CNS myelin displays sheath-to-sheath heterogeneity of its protein composition. We speculate that additional myelin proteins remain to be identified that are specific to a subset of myelin sheaths. Such proteins may have functional relevance for the local modulation of their myelin sheaths, and possibly of other cellular structures with which the latter interact. Our data do not exclude the possibility that axonal signals may modulate oligodendroglial *PMP2* expression, for example in subclusters associated with large-diameter axons (Seeker et al., 2023). On the other hand, transgenic expression of human FABP8/PMP2 in CNS myelin of mice not only affects the lipid composition of myelin but also the morphology of the axon/myelin unit, notwithstanding that the latter effect is temporary and modest. Our work emphasizes that aspects of myelin biology exist that differ between mice and humans, including the presence of FABP8/PMP2 in human CNS myelin and species-dependent differences in the lipid composition. Our findings may also be relevant for understanding the pathobiology of myelin-related disorders.

## Materials and Methods

### Human CNS samples

*Post-mortem* human optic nerve and spinal cord samples were provided by the University Medical Center Göttingen (UMG). Donors or their next of kin gave their consent to autopsy and utilization of tissue for scientific research. This was ethically approved by the respective medical ethics committees, and diagnosis was authenticated by a neuropathologist.

**Table 1.** Optic nerve tissue used for immunohistochemistry, confocal and STED microscopy, g-ratio and diameter analysis:

| Subject | Sex | Age (years) | Diagnosis | Region | Experiment | Comments |
| --- | --- | --- | --- | --- | --- | --- |
| 1 | Female | 83 | Non-diseased control | Optic nerve | IHC, STED microscopy, g-ratio and diameters | Figure 1,2 |
| 2 | Male | 81 | Non-diseased control | Optic nerve | IHC, STED microscopy, g-ratio and diameters | Figure 1,2 |
| 3 | Male | 68 | Non-diseased control | Optic nerve | IHC, STED microscopy, g-ratio and diameters | Figure 1,2 |
| 4 | Male | 59 | Non-diseased control | Optic nerve | IHC, confocal and STED microscopy, g-ratio and diameters | Figure 1,2 |
| 5 | Female | 48 | Non-diseased control | Optic nerve | IHC, STED microscopy, g-ratio and diameters | Figure 1,2 |
| 6 | Female | 66 | Non-diseased control | Optic nerve | IHC, STED microscopy, g-ratio and diameters | Figure 1,2 |
| 7 | Female | 54 | Non-diseased control | Optic nerve | IHC, STED microscopy, g-ratio and diameters | Figure 1,2 |
| 8 | Female | 54 | Non-diseased control | Optic nerve | IHC, STED microscopy, g-ratio and diameters | Figure 1,2 |

**Table 2.** Spinal cord tissue used for immunohistochemistry, confocal and STED microscopy:

| Subject | Sex | Age (years) | Diagnosis | Region | Experiment | Comments |
| --- | --- | --- | --- | --- | --- | --- |
| 6 | Female | 66 | Non-diseased control | Spinal Cord | IHC, STED microscopy | Figure 1,2 |
| 12 | Male | 64 | Non-diseased control | Spinal Cord | IHC, STED microscopy | Figure 1,2 |
| 13 | Male | 81 | Non-diseased control | Spinal Cord | IHC, STED microscopy | Figure 1,2 |
| 14 | Female | 67 | Non-diseased control | Spinal Cord | IHC, STED microscopy | Figure 1,2 |
| 5 | Male | 65 | Non-diseased control | Spinal Cord | IHC, STED microscopy | Figure 1,2 |
| 16 | Female | 84 | Non-diseased control | Spinal Cord | IHC, STED microscopy | Figure 1,2 |
| 17 | Female | 55 | Non-diseased control | Spinal Cord | IHC, STED microscopy | Figure 1,2 |
| 18 | Male | 61 | Non-diseased control | Spinal Cord | IHC, STED microscopy | Figure 1,2 |
| 19 | Female | 59 | Non-diseased control | Spinal Cord | IHC, STED microscopy | Figure 1,2 |

### Brain tissue samples from other species

Samples from brains dissected from several mammalian species were used for myelin purification. All the requirements of § 4 TierSchG, together with § 2 sentence 2, Annex 1, Section 2 and Annex 2 TierSchVersV, have been implemented. According to the German Animal Welfare Act, killing does not constitute an experiment on animals. Rat brain samples were provided by the Max Planck Institute for Multidisciplinary Sciences (Göttingen, Germany). The raccoon brain was kindly provided by a local hunter. A pig brain sample was provided by the University animal clinic (Göttingen, Germany) (notification of sacrifice for scientific purposes is documented with number T2_2022). Non-human primate brain samples were provided by the German Primate Center DPZ (Göttingen, Germany). The *post-mortem* use of samples from non-human primates with a permit in accordance with §4, §7 and §11 of the Animal Welfare Act is in line with the German legal and ethical requirements for appropriate animal experiments with non-human primates; the consultation of the Animal Welfare Committee and the Animal Welfare Officer is documented with number E2-24.

**Table 3.** Brain tissue samples from various mammalian species for myelin purification and immunoblotting:

| Species | Genotype | Region | Experiment | Comments |
| --- | --- | --- | --- | --- |
| Mouse | Control | Brain (myelin) | Myelin purification, Immunoblot analysis | Figure 1 |
| Rat | Control | Brain (myelin) | Myelin purification, Immunoblot analysis | Figure 1 |
| Raccoon | Control | Brain (myelin) | Myelin purification, Immunoblot analysis | Figure 1 |
| Marmoset | Control | Brain (myelin) | Myelin purification, Immunoblot analysis | Figure 1 |
| Rhesus Macaque | Control | Brain (myelin) | Myelin purification, Immunoblot analysis | Figure 1 |
| Baboon | Control | Brain (myelin) | Myelin purification,<br>Immunoblot analysis | Figure 1 |

### Mouse model

A new transgenic mouse line was generated under license 33.23-42502-04-21/3660 approved by the Niedersächsisches Landesamt für Verbraucherschutz und Lebensmittelsicherheit (LAVES, Oldenburg, Germany). Purified plasmid containing the CAG promoter, three SV40 polyadenylation sites flanked by loxP sites, and the *hPMP2* sequence was acquired from VectorBuilder (#VB220802–1119bhn, Chicago IL, USA). Plasmid was linearized using restriction enzymes NotI and PvuI according to the manufacturer (New England Biolabs, Ipswich, Massachusetts USA). The plasmid and enzymes were incubated together for 2.5 hours at 37°C. After enzymatic incubation, samples were diluted with a loading dye (New England Biolabs, Ipswich, Massachusetts USA) and GelRed^TM^ (Biotium, San Fransisco, USA) for UV visibility. Agarose gel (0.8%) was loaded and let run for 2 h at 130 V. Next, bands were inspected under UV for correct enzymatic activity and the relevant band was cut out with a scalpel. DNA purification was performed with the QiaQuick (Qiagen, Hilden Germany) gel purification kit, and DNA purity/concentration was measured with nanodrop 2000 (ThermoFisher, Waltham, USA). The purified DNA was diluted in injection buffer containing 5 mM Tris, 0.1 mM EDTA, pH = 7.4. Mice were generated by the transgenic facility of the MPI-NAT (Göttingen, Germany) by DNA-microinjection into embryos at the zygote-state, which were derived from C57Bl6/N mice. Subsequently, the manipulated embryos were surgically transferred into the oviduct of pseudopregnant 0.5 days post coitum (dpc) recipient-mothers using standard procedures. Founder mice were bred with C57Bl6/N mice. Next, PMP2 tg+ mice were crossbred with mice expressing *Cre* under the *Cnp* promoter (*Cnp1^tm1(cre)Kan^*), also termed *Cnp^Cre/Wt^* (Lappe-Siefke et al., 2003) yielding *PMP2^OE^;Cnp^Cre/WT^*; also termed PMP2OE mice. Experiments were performed in mutant mice (PMP2OE) and respective controls (WT/WT, *Cnp^Cre/WT^* or *PMP2^OE^;Cnp^WT/WT^*). Genotyping was performed by genomic PCR. *PMP2* tg+ mice were identified using sense primer 5′-AGCTTATCGATACCGTCACCCAGC in combination with antisense primer 5′-GCACTTGATTCAGTGATCCTCTCTGC yielding a 333 bp product. Control mice had no visible band. For the recombination PCR, sense primer 5′-AGAACTAGTGGATCCGGAACCCTT in combination with the same antisense primer was used to detect recombination yielding a 387 bp fragment. To detect the *Cnp^Cre^* allele (Lappe-Siefke et al., 2003) we used the same primers as in (Buscham et al., 2022).

### Animal welfare

Mice were bred and raised at the MPI-NAT (Göttingen, Germany) with 2-4 mice per cage, had access to food (Ssniff R/M Haltung V1534-000, Ssniff, Soest, Germany) and water *ad libitum,* and experienced a 12-hour light-dark cycle. All experiments were in accordance with the German Animal Welfare Law (Tierschutzgesetz der Bundesrepublik Deutschland, TierSchG). For procedures of sacrificing rodents for subsequent preparation of tissue, all regulations given in TierSchG §4 were followed. Since sacrificing of mice is not an experiment on animals according to TierSchG §7 Abs. 2 Satz 3, no specific authorization was required. The animal facility at the MPI-NAT is registered at the Niedersächsisches Landesamt für Verbraucherschutz und Lebensmittelsicherheit (LAVES) according to TierSchG §11 Abs. 1. According to TierSchG and the regulation about animals used in experiments dated 11^th^ August 2021 (Tierschutz-Versuchstierverordnung, TierSchVersV), an animal welfare officer and an animal welfare committee are established for the institute.

### Myelin purification

Myelin was purified from nervous tissue using an established protocol including two sucrose gradient ultracentrifugation steps and two osmotic shocks (Erwig et al., 2019). Myelin accumulates as light weight fraction between the sucrose concentrations (0.32 M and 0.85 M).

### Immunoblotting

Protein concentration measurements and gel electrophoresis was conducted as in (Gargareta et al., 2022). Immunoblotting was performed as in (Patzig et al., 2016, Gargareta et al., 2022). Primary antibodies were specific for SEPTIN7 (IBL, #IBL18991, 1:1000), SEPTIN8 (ProteinTech, #11769-1-AP, 1:500), PLP/DM20 (A431 (Jung et al., 1996), 1:5000), FABP8/PMP2 (ProteinTech, #12717-1-AP, 1:500), 2’,3’-Cyclic nucleotide 3’-phosphodiesterase (CNP, Abcam, #C5922, 1:1000), MOG (8-18C5 (Linnington et al., 1984), 1:500), sirtuin 2 (SIRT2, Abcam, #ab67299, 1:500), glial fibrillary acidic protein (GFAP, DAKO, #Z0334, 1:1000), and Class III beta-tubulin (TUJ1, Biolegend, #MMS-435P, 1:500). Appropriate secondary anti-mouse, anti-rabbit and anti-rat antibodies conjugated to HRP were from Dianova (HRP goat-anti-mouse, #115-035-003, 1:5000; HRP goat-anti-rabbit, #111-035-003, 1:5000; HRP goat-anti-rat, #112-035-167, 1:5000). Fast Green staining of membranes was performed as detailed in (Gargareta et al., 2022). Immunoblots were developed using the Enhanced Chemiluminescence Detection kit (Western Lightning Plus, Perkin Elmer, Waltham, MA) and the Super Signal West Femto Maximum Sensitivity Substrate (Thermo Fisher Scientific, Rockford, IL). Signal was detected using the Intas ChemoCam system (INTAS Science Imaging Instruments GmbH, Göttingen, Germany).

### Myelin proteome analysis

Myelin proteins were digested in-solution using filter-aided sample preparation (FASP) as described (Erwig et al., 2019) and subsequently analyzed by LC-MS employing different MS^E^-type data-independent acquisition (DIA) strategies, as recently established for mouse PNS and CNS myelin (Jahn et al., 2020; Siems et al., 2020). Protein fractions equivalent to 10 μg of myelin protein were solubilized in reducing lysis buffer (1% ASB-14, 7 M urea, 2 M thiourea, 10 mM DTT, 0.1 M Tris, pH 8.5) and alkylated with iodoacetamide. Samples were processed following a CHAPS-based FASP protocol using centrifugal filter units (30 kDa MWCO, Merck Millipore) and after detergent removal, the urea buffer was exchanged for digestion buffer (50 mM ammonium bicarbonate (ABC), 10% acetonitrile). Proteins were then digested overnight at 37 °C with 400 ng trypsin. Tryptic peptides were collected by centrifugation and subsequently extracted with 40 µl of 50 mM ABC and 40 µl of 1% trifluoroacetic acid (TFA). For quantification according to the TOP3 approach, combined flow-throughs were spiked with 10 fmol/μl of Hi3 EColi standard (Waters Corporation; contains a set of quantified synthetic peptides derived from *E. coli.* chaperone protein ClpB) and directly subjected to LC-MS-analysis. The samples were first analyzed in the ion mobility-enhanced DIA mode with drift time-specific collision energies referred to as UDMS^E^ (providing increased proteome coverage at the cost of dynamic range for the accurate quantification of medium- to low-abundant proteins) and rerun in a data acquisition mode without ion mobility separation of peptides referred to as MS^E^ (providing increased dynamic range at the cost of proteome coverage for the accurate quantification of exceptionally abundant myelin proteins) (see (Siems et al., 2020) for details). Continuum LC-MS data were processed using Waters ProteinLynx Global Server (PLGS) and searched against a customized database comprising the UniProtKB/Swiss-Prot mouse proteome (release 2024-01, 17,201 entries), supplemented with sequence data for E. coli chaperone protein ClpB, porcine trypsin, and human PMP2. Reversed sequences were included to enable false discovery rate (FDR) estimation, which was controlled at 1%. Post-identification analysis, including TOP3-based protein quantification, was carried out using the freely available software ISOQuant (Distler et al., 2014), as previously described (Jahn et al., 2020; Siems et al., 2020). Proteins were quantified as parts per million (PPM), i.e., the relative abundance (w/w) of each protein with respect to the total protein content, but only if they were represented by at least two peptides (minimum length of six amino acids, score ≥ 5.5) identified in at least two runs. A false discovery rate (FDR) threshold of 1% was applied at both the peptide and protein levels, and at least one unique peptide was required. Known contaminants from blood (albumin, hemoglobin) and skin or hair (keratins) were excluded. Potential outliers were further assessed by examining peptide identification quality, quantification consistency, and distribution across isoforms. Filtered protein datasets were subjected to statistical analysis using the Bioconductor R packages *limma* and *q-value* to detect significant differences in protein abundance via moderated t-statistics, as previously outlined (Jahn et al., 2020; Siems et al., 2020). Proteome profiling of control and PMP2OE myelin included three to five biological replicates per genotype, each analyzed in duplicate digestions as technical replicates. For visualization, volcano plots were generated in GraphPad Prism 10 by plotting q-values against fold changes after −log10 and log2 transformation, respectively. Color coded heatmaps were prepared in Microsoft Excel 16.93.

### Myelin lipidome analysis

Biochemically purified myelin (see above) was mixed 1:1 with methanol. Aliquots of 130 µl were mixed with 195 µl ultrapure water and 650 µl 1-butanol and vortexed for 3 min. Subsequently, phase separation was assisted by centrifugation for 5 min at 20000 g. The organic phase was transferred to a 1.6 ml glass vial and 325 µl of water-saturated butanol was added to the aqueous phase and vortexed for 3 min. After centrifugation for 5 min at 20000 g the organic phase was added to the organic phase of the first extraction and the combined organic phase was dried in a speedvac. Dry lipds were resuspended in 40 µL of a 2:1:1 mixture of 2-propanol, acetonitrile, and water.

Non-targeted measurement of lipids by LC-MS was carried out as described previously (Edwards-Hicks et al., 2020) using an Agilent 1290 Infinity II UHPLC inline with a Bruker Impact II QTOF-MS operating in negative or positive ion mode. Briefly, the scan range was from 50 to 1600 Da. Mass calibration was performed at the beginning of each run. LC separation was on a Zorbax Eclipse plus C18 column (100 x 2 mm, 1.8 µm particles) using a solvent gradient of 70% buffer A (10 mM ammonium formiate in 60:40 acetonitrile:water) to 97% buffer B (10 mM ammonium formiate in 90:10 2-propanol:acetonitrile). Flow rate was 400 µl/min, autosampler temperature was 5 °C, and injection volume was 2 µl. Data processing including feature detection, feature deconvolution, and annotation of lipids was performed using MetaboScape (version 2025). Filtering for artifacts and outliers was performed in R (see **Supplemental Data Table S3**).

### PMP2 immunolabelling on human oligodendrocytes transplanted into mouse brain

PMP2 immunostaining on human-mice transplant tissue was performed using previously generated and published material (Ramesh et al., 2025). In brief, human OPCs derived from control human embryonic stem cells (hESCs) were transplanted into hypomyelinated and immunosuppressed newborn mouse pups (*Mbp^shi/shi^; Rag1^-/-^)* and analyzed at 11-12 weeks post transplantation. Cryosections of the transplanted brains were generated as described (Ramesh et al., 2025). For PMP2 immunohistochemistry, slides containing cryosections were briefly washed in 1x PBS (Gibco™ PBS, pH7.4, 70011044, Life Technologies LTD, Paisley, UK) before permeabilizing for 15 min with 0.5% Triton X-100 (Merck Life Sciences UK Limited, X100-1L, Glasgow, UK) in 1x PBS. This was followed by blocking with 5% normal goat serum (S1000, Vector labs, Newark, CA, USA), 0.1% Triton X-100 in 1x PBS for 1 h at room temperature. PMP2 and MBP primary antibodies were diluted in the same blocking solution and sections were incubated for 16-20 h at 4°C. Sections were washed in 1x PBS (3×15 min each) and incubated with secondary antibodies for 1 h at room temperature. Following 3x 15 min washes with 1x PBS, the slides were counterstained with Hoechst (62249, Thermo Fisher, Scientific, Paisley, UK) for visualization of nuclei. All slides were mounted using Fluorsave mounting medium (345789, Merck Life Sciences UK Limited, Glasgow, UK) and using 24×50 mm glass coverslips. The sections were imaged using 63x objective on Zeiss LSM710 confocal microscope with 1024×1024 resolution (16 μm-thick sections, step size 0.5 μm). Corpora callosa and fornix were imaged with 2-3 FOV per section. From each mouse, 2-3 sections were analyzed. Primary antibodies were specific for PMP2 (rabbit, Proteintech, 12717-1-AP, 1:500) and MBP (rat monoclonal, Abcam, Cambridge, UK, AB7349, 1:1000). Secondary antibodies were goat anti-rat 647 (A21247, Thermo Fisher Scientific, Paisley, UK 1:1000) and goat anti-rabbit 555 (A21428, Thermo Fisher Scientific, Paisley, UK 1:1000).

PMP2 immunolabelling was also performed in brain slices from mice in which forebrain myelin was replaced by human myelin. Briefly, human iPSC-derived OPCs expressing GFP were transplanted into immunodeficient mouse pups (P1) in which normal myelination of the forebrain is prevented (genotype *Myrf^flox/flox^;FoxG1^Cre+/WT^;Rag2^-/-^*). Mice were aged for 6 months to allow human oligodendrocyte differentiation and myelination of the forebrain. Following transcardial perfusion-fixation with 4% paraformaldehyde, 100 μm brain sections were cut using a vibratome. Slices were incubated in permeabilization and blocking solution (10% goat serum, 0.5% Triton X-100 in PBS) for 4-6 hours. Slices were incubated with primary antibodies overnight at room temperature. After washing, slices were incubated with secondary antibodies in PBS overnight at 4 °C. After washing, slices were stained with DAPI (Sigma-Aldrich, D9542) for cell nuclei and then mounted (Fluoromount-G). Primary antibodies used were specific for PMP2 (rabbit, Proteintech 12717-1-AP, 1:200), MBP (rat, Biorad #2287, 1:50) and GFP (chicken, Abcam, ab1370, 1:1000). Secondary antibodies were goat anti-chicken Alexa 488 (Abcam ab150169), goat anti-rat 568 (Thermo Fisher A11077) and goat anti-rabbit Alexa 647 (Life Technologies A21245).

### Immunohistochemistry

For immunolabelling of nervous tissue, paraffin-embedded non-diseased human optic nerve, spinal cord, and brain, as well as mouse sciatic nerve and spinal cord were processed as described (Gargareta et al., 2022). After rinsing samples with 0.05 M Tris buffer (pH 7.6) containing 2% milk powder (Frema Instant Magermilchpulver, granoVita, Radolfzell, Germany) for 1 × 5 min all sections were put into cover plates (besides non-diseased human optic nerves, which were instead covered with parafilm) and then blocked with 10% goat serum (Gibco/Thermo Fisher Scientific #16210064, Waltham, MA) diluted 1:4 in PBS (pH 7.4)/ 1% BSA. Finally, primary antibodies diluted in 1x PBS were applied overnight at 4°C. Next, sections were washed 3 × 5 min with Tris buffer and secondary antibodies or secondary nanobodies were applied in incubation buffer (1x PBS) for 2 hours. Slides were then rinsed again 3 × 5 min with Tris buffer and DAPI was applied in concentration of 1:2000 for 5 min. Finally, slides were again washed briefly 2 × 5 min with Tris buffer and sections were mounted using Aqua-Poly/Mount (Polysciences, Eppelheim, Germany). Antibodies were specific for Proteolipid protein 1 (PLP1/DM20 (A431), kind gift from Martin Jung, 1:200), Peripheral Myelin Protein 2 (PMP2, Proteintech, #12717-1-AP, 1:100-1:400) and Myelin basic protein (MBP, Serotec, #MCA4085, 1:200-1:400). Secondary antibodies were donkey anti-rat Alexa 488 (Invitrogen, #1979698, 1:400) and goat anti-rat DyLight 488 (Invitrogen, #SA510018, 1:400). Secondary nanobodies were FluoTag®-X4 anti-rabbit IgG Aberrior Star635P (NanoTag, #N2404, 100-120nM).

### Microscopy

#### STED microscopy

STED imaging of non-diseased optic nerves and spinal cords was performed using a STED microscope (Abberior, Göttingen, Germany), which consisted of an IX83 inverted microscope (Olympus, Hamburg, Germany) equipped with an UPLSAPO 100x 1.4 NA oil immersion objective (Olympus). An Autofocus system (Abberior, Göttingen, Germany) was pre-installed to compensate for z-drifts during image acquisition. For dual channel imaging, excitation lasers of 640 nm and 488 nm, and depletion lasers of 775 nm and 595 nm were adjusted to ∼2 μW and ∼10 mW, respectively, as measured using a microscope slide power sensor (S170C, Thorlabs, Inc.) and a power and energy meter interface (PM100USB, Thorlabs, Inc.). The (x-y) pixel size was set to 20 nm, and the dwelling time was set to 10 μs.

For g-ratio analysis, 20 randomly selected, distinct images were acquired after IHC at 100x magnification using STED microscopy; ImageJ (Fiji) was used for g-ratio assessment. The g-ratio was calculated as the ratio between the axonal diameter of the circle of equal area and the diameter of the circle of equal area of the corresponding myelin sheath (Fledrich et al., 2014; Patzig et al., 2016). For the analysis only normal-appearing myelinated axons were used and were further categorized as PMP2-immunopositive or PMP2-immunonegative. In total, between 35 and 160 axons per non-diseased human optic nerve with n=8 were quantified. The same procedure was followed for the non-diseased spinal cord. There, we measured 37-171 axons in n=9 spinal cords.

Additional images from the non-diseased human spinal cord were acquired using a STEDYCON microscope (Abberior, Göttingen, Germany). The microscope consisted of an IX83 inverted microscope (Olympus, Hamburg, Germany) equipped with a 100x/1.45 UPLXAPO100XO, NA: 1.45, oil immersion objective (Olympus) and a Stedyfocus system (Abberior, Göttingen, Germany). For double-channel imaging, we used excitation lasers at 488 nm and 640 nm, and a depletion laser at 775 nm. The (x-y) pixel size was set to 25 nm, and the dwelling time was set to 10 μs.

#### Confocal microscopy

Confocal microscopy of non-diseased human optic nerves, mouse sciatic nerve and humanized mouse brain was performed using the confocal microscope LSM880 (Zeiss, Oberkochen, Germany) containing Plan-Apochromat 40x/1.4 Oil DIC M27 and Colibri (Zeiss, Oberkochen, Germany) 5 LED light source. A FS90 filter was used to observe the samples with the light source Colibri. DAPI was excited at 405, Alexa 488 or Dylight 488 was excited with an Argon Laser at an excitation of 488 and Abberior635P was excited with a HeNe633 laser at 633 nm. Finally, the MBS 488/561/633 beam splitter was used to detect Alexa 488 and Abberrior635P and MBS-405 for DAPI respectively.

We additionally used the STEDYCON (Abberior, Göttingen, Germany) microscope for confocal imaging acquisition of mouse brain and spinal cord as well as for non-diseased human spinal cord and humanized mouse brains. The microscope consists of an IX83 inverted microscope (Olympus, Hamburg, Germany) equipped with an 100x/1.45 UPLXAPO100XO, NA: 1.45, oil immersion objective (Olympus) and a Stedyfocus system (Abberior, Göttingen, Germany). For triple channel imaging, excitation lasers of 405 nm, 488 nm, and 640 nm were used. The (x-y) pixel size was set to 100 nm and dwelling time to 3 μs.

All images were analyzed with Fiji. Brightness and contrast intensity of channels was adjusted accordingly, and background staining was subtracted using a rolling ball radius of 30-50 pixels. Further analysis was performed using the Gaussian Blur filter and a sigma (radius) of 0.50-1. Adobe Illustrator was used to export, process and assemble the figures.

### Electron Microscopy

#### TEM microscopy

Conventional transmission electron microscopy (TEM) of mouse optic nerves was performed as in (Eichel et al., 2020; Gargareta et al., 2024; Weil et al., 2019). Briefly, optic nerves of PMP2OE mice and respective controls were dissected at P24 (n=5) and P40-45 (n=4) and fixed in fixative (4% PFA, 2.5% glutaraldehyde in 0.1 M phosphate buffer) at room temperature and stored at 4°C until further processing. Thereafter, optic nerves were postfixed with OsO_4_ (Science Services, Munich, Germany), dehydrated with acetone and embedded in epoxy resin (Serva). For TEM, ultrathin sections (50 nm) were cut with the use of a diamond knife (45, Diatome Biel, CH) via the PTPC Powertome Ultramicrotom (RMC, Tuscon Arizona, USA) and collected on formvar-coated hexagonal 100 mesh copper grids (Science Services, Munich, Germany). UranyLess (Electron Microscopy Science, Hatfield, Panama) was used to contrast ultrathin sections for 15–30 min followed by 6 x 5 min subsequent washing steps with ddH2O (Patzig et al., 2016; Weil et al., 2019). Samples were examined using EM 912 AB-Omega (Zeiss, Oberkochen, Germany) coupled to a 2k CCD camera (TRS, Moorenweis, Germany). For quantitative analyses of g-ratio, axonal diameters, normal and pathological appearing profiles as well as myelinated and unmyelinated axons at P24 and P40, 10 random, non-overlapping images were taken at 7000x and 4000x magnification respectively. All analyses were assessed using ImageJ (Fiji). The g-ratio was calculated as the ratio between the axonal diameter of the circle of equal area and the diameter of the circle of equal area of the corresponding myelin sheath (Fledrich et al., 2014; Patzig et al., 2016). For g-ratios only normal-appearing, non-degenerating/degenerated myelinated axons were analyzed. All quantifications were performed blinded to the genotype.

Further axonal diameter analysis of 10 different and independent EM images at P40-P45 (n-4) was performed using the Myeltracer software. The software generates a greyscale image and applies a manually adjusted threshold based on black and white pixels thus defining the axon area (Kaiser et al., 2021). Depending on image quality, some axons require manual tracing. Again, all quantifications were performed blinded to the genotype.

### Data processing for single-cell chromatin accessibility and mRNA abundance of *PMP2/Pmp2*

Expression and chromatin accessibility of *PMP2*/*Pmp2* and a panel of selected genes was obtained from published single cells multiome datasets from human prefrontal cortex and spinal cord (Kabbe et al., 2026), mouse spinal cord (Zheng et al., 2025), and mouse brain(Bravo González-Blas et al., 2023). For human, OPC and MOL annotated cells were subset from the original published annotations (Kabbe et al., 2026). For mouse spinal cord, the original published multiome dataset was subset to retrieve only CFA control samples and as annotations only cells assigned as OPC and mature MOLs were selected. To keep number of cells comparable between samples the mouse spinal cord dataset was subsampled randomly to 500 cells for each OPCs and MOLs. The genome assemblies used are GRCm38-mm10 for *Mus musculus* and GRCh38-hg38 for *Homo sapiens* data. For the mouse brain, the raw FastQ files from SRA (SRR20963904 and SRR20963906) with GEO accession number GSE210747 were downloaded and processed. Reads were aligned with cellranger-arc.2.0.2 to mouse mm10 genome assembly (refdata-cellranger-arc-mm10-2020-A-2.0.0). Gene expression and chromatin accessibility matrixes were processed with Seurat v 5.3.0 (Hao et al., 2024) and Signac v 1.14.0 (Stuart et al., 2021), cells were filtered with following parameters; nCount_RNA < 25000 & nCount_RNA > 200 & nucleosome_signal < 2 & TSS.enrichment > 1 & percent.mt < 20. For each modality, RNA and ATAC, dimension reduction was performed in each individually (PCA 1:50, LSI 2:50) by getting each modalities weights and constructing the Weighted Nearest Neighbor (WNN) graph with both from Seurat. Gene activities from chromatin accessibility were calculated using 500 bp upstream the annotated transcription start sites plus the full genebody. Gene activity was only recalculated for mouse brain scATAC-seq. To identify MOL and OPCs label transfer, Seurat FindTransferAnchors (CCA) on the intersecting variable genes between both datasets, and manual checking of canonical markers was used with the reference mouse atlas (Zeisel et al., 2018). Dotplots were generated by extracting the $data slot from the DotPlot Seurat function for each of the datasets for the selected genes (*PMP2*/*Pmp2*, *TSPAN2*/*Tspan2*, *SOX10*/*Sox10*, *MYRF*/*Myrf, PDGFRA*/*Pdgfra*). In the RNA assay for expression and gene activity assay for chromatin accessibility, final dotplots show the log2 absolute average expression of each gene in each cluster on the plot; scales are comparable within the plot. Chromatin accessibility BigWig tracks for OPCs and MOLs were constructed by selecting the fragments with barcodes annotated as OPC or MOL in each dataset. OPC and MOL fragments were converted to bam files, BedTools v.2.25.0 (Quinlan and Hall, 2010) bedtobam and Samtools v1.16.1 (Li et al., 2009). Bams were converted to BigWig with Deeptools v3.4.3 (Ramírez et al., 2016) bamCoverage --normalizeUsing RPKM --binSize 50 --centerReads --smoothLength 250 --extendReads 200. Browser capture was built with WashU Epigenome Browser (Li et al., 2019; Zhuo et al., 2023). The code will be made available at https://github.com/Castelo-Brancolab/PMP2_Pmp2_manuscript.

### Statistics

Statistical analyses were mainly performed in GraphPad Prism 10 (GraphPad Software, Inc., San Diego, United States) or R-studio. Data in bar graphs show mean ±SD (error bars), except for diameters in **Figures 2D,2I**, and **5K**, and **Supplemental Figure S4H**, which show median ±SD (error bars). Data points in bar graphs represent individual human subjects (**Figure 2, supplemental Figure S2)** and individual mice **(Figure 5, supplemental Figure S5).** Data points in data clouds showing g-ratios represent individual axon/myelin profiles in **Figure 2C,H, 5J, supplemental Figure 5H**. Size/number is provided in figures or figure legends. Two-tailed Student’s t-test was applied to compare two groups. Relative frequency distribution of axonal diameters was assessed statistically by RStudio (www.rstudio.com, Version 3.4.1; https://github.com/JeneaBin/AxonDiameterStats/releases/tag/v1.0.0) as described (Bin et al., 2025), and visualized using GraphPad Prism 10.

Lipid levels **(Figure 6B,C)** were compared between groups using unpaired two-tailed *t*-tests, assuming Gaussian distribution. To account for multiple comparisons, adjusted *p*-values were calculated using the Bonferroni–Šidák correction (family-wise error rate control). In all tests, significance was set at p-value of <0.05. Significance levels are depicted as n.s.=non-significant, *p<0.05, **p<0.01, and ***p<0.001 with p-values given in respective figure legends. Representative STED micrographs were chosen from the images obtained from eight different human optic nerves and nine human spinal cords. STED microscopy representative images were chosen from three different independent experiments. Representative electron micrographs were choses from the images obtained from n=5 control and PMP2OE mice at P24 and n=4 at P45. All analyses were performed blinded to the genotype.

## Data availability statement

All relevant data are included in the main paper or supplemental files. The mass spectrometry proteomics data have been deposited to the ProteomeXchange Consortium via the PRIDE (Perez-Riverol et al., 2025) partner repository with dataset identifier PXD071876. The mass spectrometry lipidomics data have been deposited to the MassIVE repository with dataset identifier MSV000102070. Datasets will be accessible upon publication in a scientific journal.

## Acknowledgements

We thank U. Bode, A. Bretschneider, A. Fahrenholz, L. Piepkorn, T. Ruhwedel, B. Sadowski, M. Schindler, and K. Wilke for technical support. We thank C. Schuon for the pig brain sample, K. Mätz-Rensing for non-human primate brain samples, L. Linhoff for the rat brain sample, and a local hunter for the raccoon brain sample. We thank I. Huitinga, U. Fünfschilling, S. Kimmina, T. Papadopoulos, and A. Schräpler for discussions and support. We thank the transgenic facility at the MPI-NAT City Campus headed by T. Papadopoulos for transgenesis and A. Diedrich, S. Lempa, and M. Peine for mouse husbandry. We thank C. Linington and M. Jung for antibodies. V.-I.G. and S.B.S. thank the International Max Planck Research School for Genome Science (IMPRS-GS) for support. This work was supported by the Deutsche Forschungsgemeinschaft (DFG grant WE 2720/5-1 to H.B.W.), and the Adelson Program for Neurodegenerative Diseases (APND to K.-A.N.). The project was in part funded by the Federal Ministry of Education and Research (Bundesministerium für Bildung und Forschung, BMBF) as part of the German Center for Child and Adolescent Health (DZKJ, funding code 01GL2402A to C.S.), the SFB 274/2 (project ID 408885537, project B03 to C.S.), and by the Research Council of Norway (BioPROM project, project number 324877 to P.K.). Open access publication was funded and organized by project DEAL.

## Author contributions

VIG Investigation, Data analysis, Writing-original draft, Writing-review & editing

NM Investigation, Writing-review & editing

ETH Investigation, Writing-review & editing

SBS Methodology, Writing-review & editing

SJC Investigation, Writing-review & editing

BV Investigation, Writing-review & editing

RTK Unpublished materials, Supervision, Writing-review & editing

RBJ Investigation, Writing-review & editing

JB Investigation, Writing-review & editing

VR Investigation, Writing-review & editing

BTS Supervision, Writing-review & editing

SC Supervision, Writing-review & editing

LZ Unpublished materials, Supervision, Writing-review & editing

JMB Investigation, Data analysis, Writing-review & editing

MAEV Methodology, Writing-review & editing

DAL Supervision, Writing-review & editing

EA Investigation, Writing-review & editing

TS Investigation, Writing-review & editing

GCB Supervision, Writing-review & editing

MU Investigation, Writing-review & editing

LvW Investigation, Writing-review & editing

SZ Investigation, Writing-review & editing

UF Investigation, Writing-review & editing

AF Supervision, Writing-review & editing

WM Methodology, Writing-review & editing

RS Unpublished materials, Writing-review & editing

AJL Investigation, Writing-review & editing

PK Supervision, Writing-review & editing

CS Unpublished materials, Writing-review & editing

OJ Investigation, Methodology, Data analysis, Writing-review & editing

KAN Supervision, Funding, Writing-review & editing

FO Supervision, Data analysis, Writing-review & editing

HBW Conceptualization, Supervision, Data analysis, Funding, Writing-original draft, Writing-review & editing

---

**Supplemental Figure S1.**
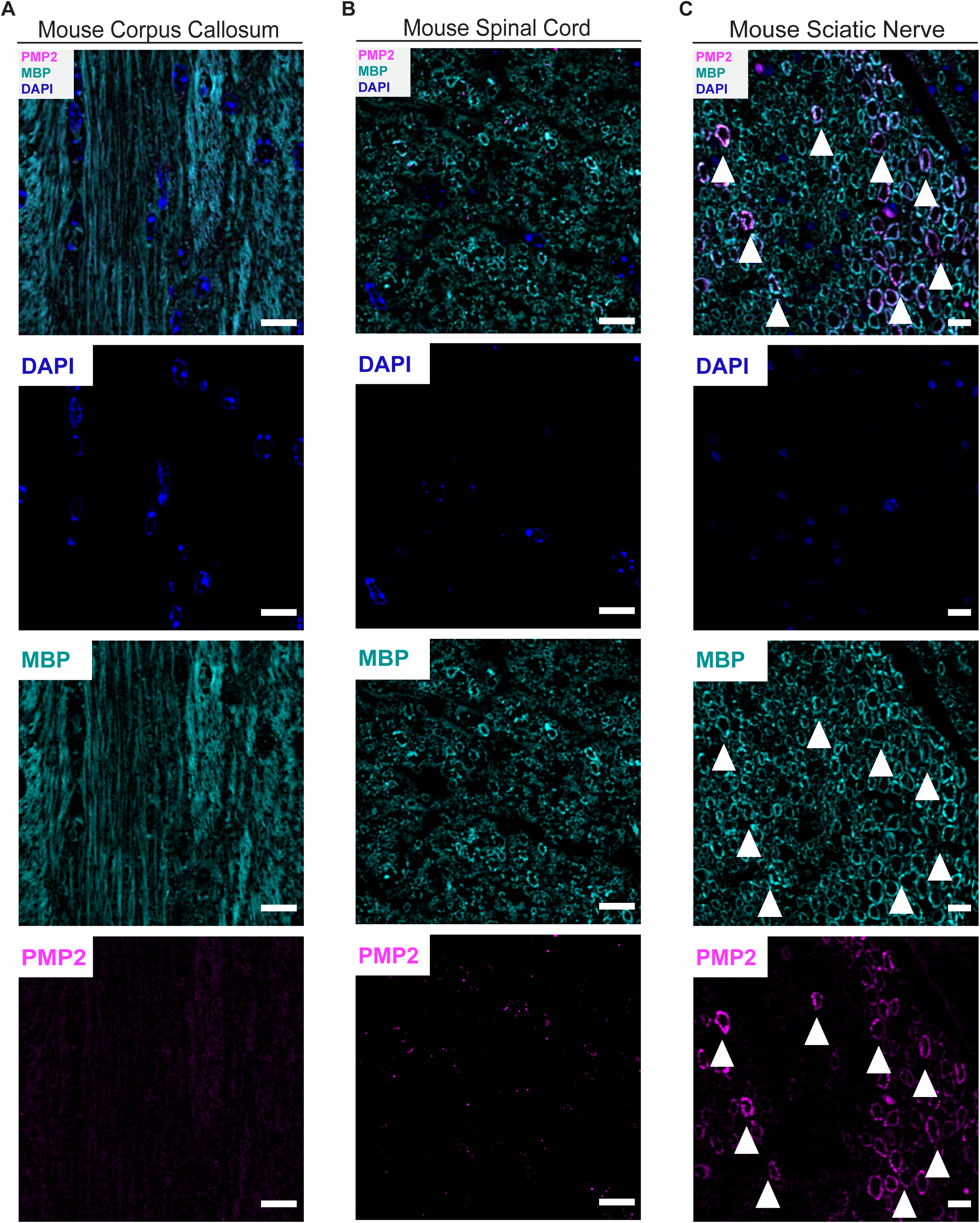
FABP8/PMP2 is expressed in PNS but not CNS myelin of mice. **A,B** Immunofluorescence labelling and confocal microscopy detecting PMP2 (magenta) and MBP (cyan) on cross-sectioned corpora callosa **(A)**, spinal cords **(B),** and sciatic nerves **(C)** of C67Bl/6N mice at 9 mo. DAPI in blue. Arrowheads point at myelin rings double-immunopositive for MBP and PMP2. Note that in mice PMP2 immunolabeling was readily detected in a subset of myelin sheaths and Schwann cells in sciatic nerves that are part of the PNS, in agreement with previous reports (Trapp et al., 1979; Yim et al., 2022), but not in spinal cords and corpora callosa of mice that are part of the CNS. Scale bar 10 μm.

**Supplemental Figure S2.**
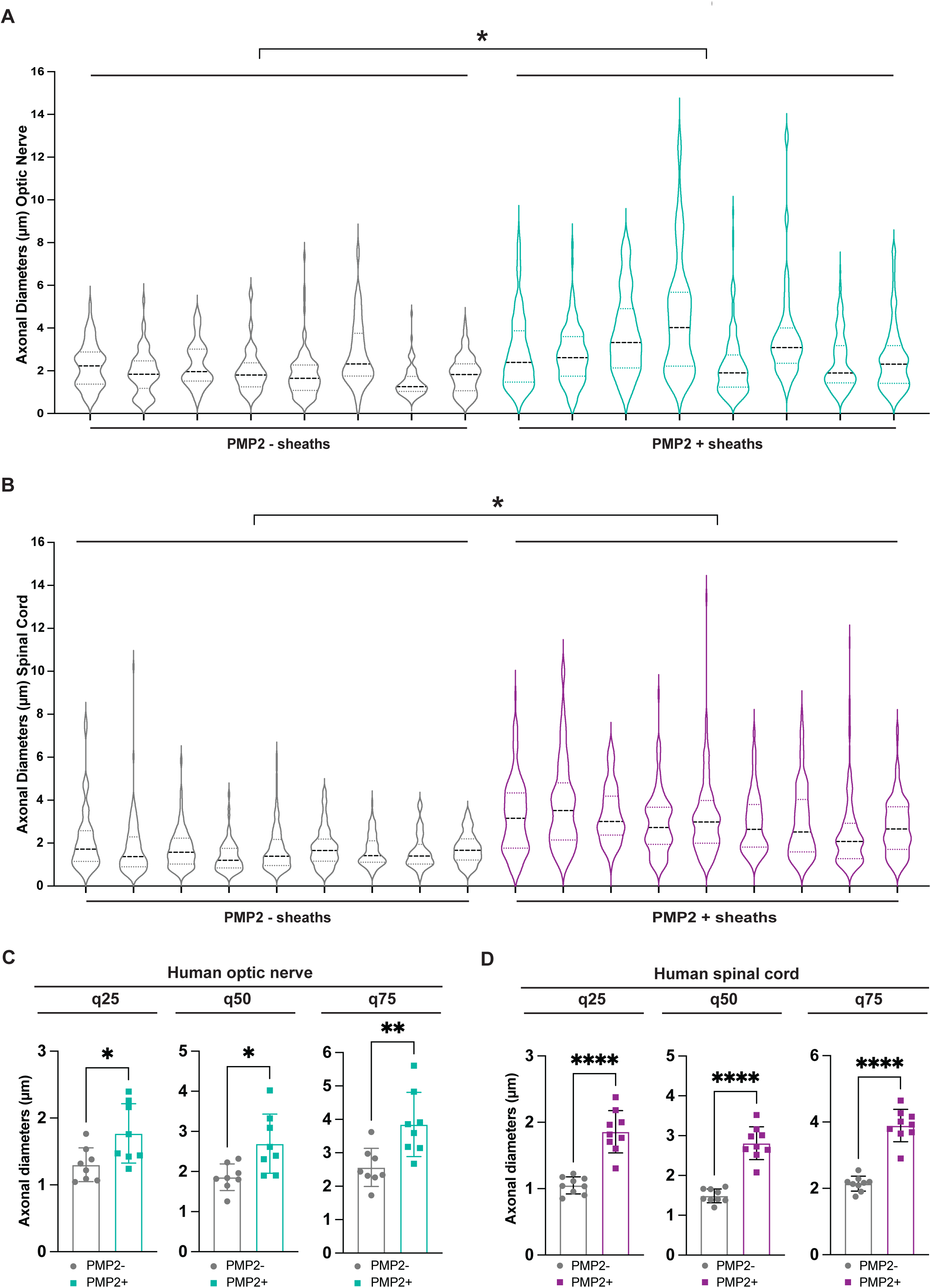
Analysis of diameters of human CNS axons with PMP-immunopositive versus PMP-immunonegative myelin sheaths. **A** Violin plot showing diameters of axons with PMP2-immunonegative (PMP2-, gray) and axons PMP2-immunopositive (PMP2+, turquois) myelin sheaths in human optic nerves as assessed on images as in **Figure 2A**. Each violin represents one out of eight human individuals; horizontal lines indicate the 25th, 50th (median), and 75th percentiles to show spread and skewness. Evaluation of the distribution of the diameters using quantile analysis (lqmm) identified differences in axon diameters at the quartile boundaries (first and second quartile boundary: n.s p = 0.1148; median: ** p = 0.002818; third and fourth quartile boundary: *p = 0.01419). **B** Violin plot showing diameters of axons with PMP2-immunonegative (PMP2-, gray) and axons PMP2-immunopositive (PMP2+, magenta) myelin sheaths in human spinal cord as assessed on images as in **Figure 2B**. Each violin represents one out of nine human individuals; Horizontal lines indicate the 25th, 50th (median), and 75th percentiles to show spread and skewness. Evaluation of the distribution of the diameters using quantile analysis (lqmm) identified differences in axon diameters at the quartile boundaries (first and second quartile boundary: ** p = 0.003386; median: *** p < 0.001; third and fourth quartile boundary: ***p < 0.001). **C** Bar graphs showing comparison of means across quartile groups (q25, 25%; q50, 50%; q75, 75%) of axon diameters in n=8 human optic nerves comparing axons myelinated by PMP2-immunonegative (PMP2-, gray) or PMP2-immunopositive (PMP2+, turquoise) myelin sheaths in the optic nerves of humans. Each datapoint represents one individual. Bar graph gives mean ±SD; q25 * p=0.0215, q50 * p=0.0113, q75 ** p=0.0057 by two-tailed Student’s t-test. Note that the mean diameters of axons myelinated by PMP2 immunonegative and PMP2-immunopositive myelin sheaths differ significantly in all quartiles. **D** Bar graphs showing comparison of means across quartile groups (q25, 25%; q50, 50%; q75, 75%) of axon diameters in n=9 human spinal cords comparing axons myelinated by PMP2-immunonegative (PMP2-, gray) or PMP2-immunopositive (PMP2+, magenta) myelin sheaths in the spinal cord of humans. Each datapoint represents one individual. Bar graph gives mean ±SD; q25 **** p<0.0001, q50 **** p<0.0001, q75 **** p<0.0001by two-tailed Student’s t-test. Note that the mean diameters of axons myelinated by PMP2-immunonegative and PMP2-immunopositive myelin sheaths differ significantly in all quartiles.

**Supplemental Figure S3.**
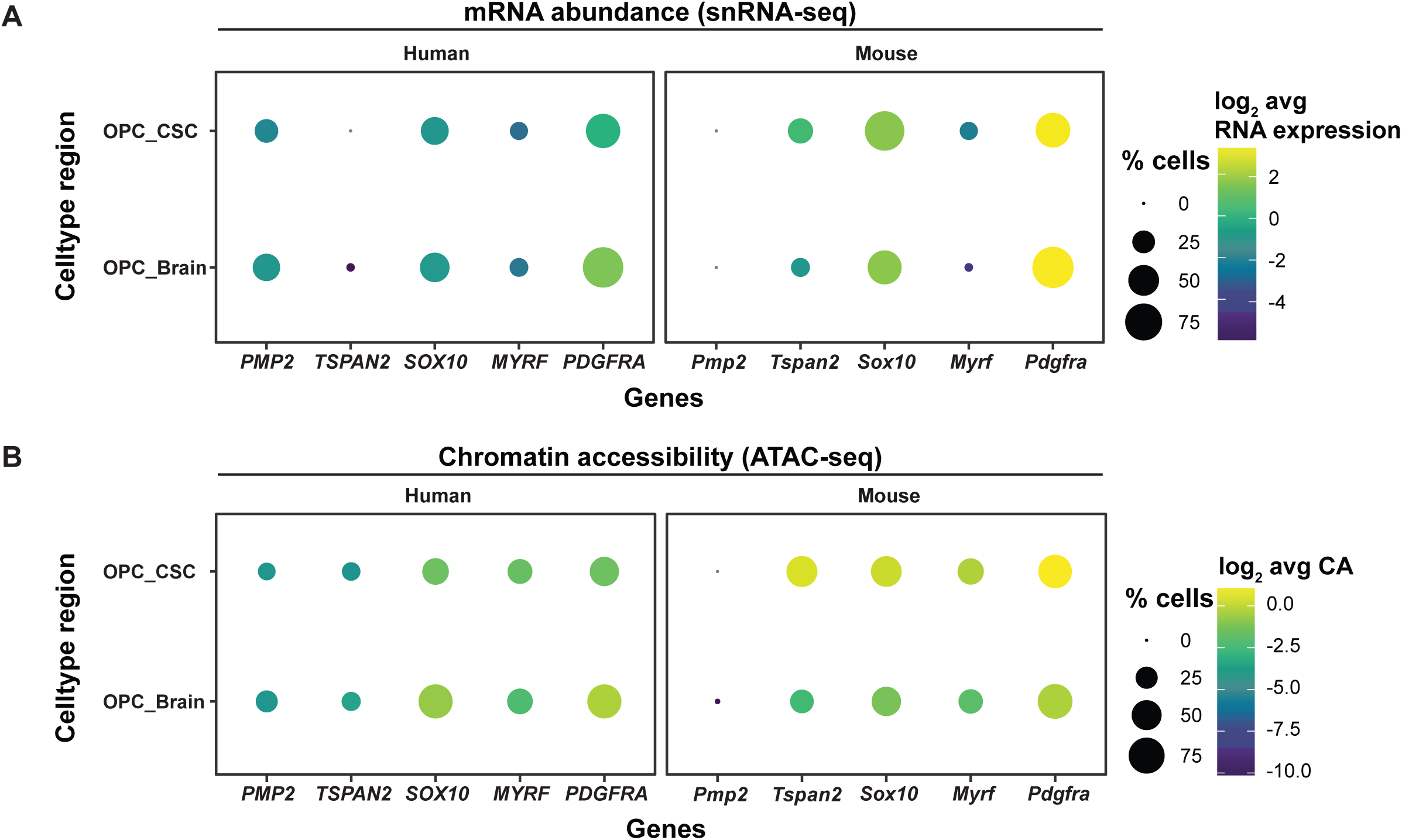
Cross-species snRNA-seq and scATAC-seq analysis of *PMP2/Pmp2* in oligodendrocyte precursor cells. **A** Dot plot showing the percentage of oligodendrocyte precursor cells (OPC) expressing *PMP2/Pmp2* and the indicated marker mRNAs, and their log_2_-transformed average mRNA abundance, in the cervical spinal cord (CSC) and brains. Data are taken from previously established human (left) and mouse (right) snRNA-seq datasets (Bravo González-Blas et al., 2023; Kabbe et al., 2026; Zheng et al., 2025). Note that *PMP2/Pmp2* is expressed in a considerable percentage of human but not mouse OPC. For myelinating oligodendrocytes (MOL) see **Figure 3A**. **B** Dot plot showing the percentage of myelinating oligodendrocytes (MOL) in which the gene locus of *PMP2/Pmp2* and the indicated marker genes is accessible, and their log_2_-transformed average chromatin accessibility, in the cervical spinal cord (CSC) and brains. Data are taken from previously established human (left) and mouse (right) scATAC-seq datasets (Bravo González-Blas et al., 2023; Kabbe et al., 2026; Zheng et al., 2025). Note that the *PMP2/Pmp2* locus is accessible in a considerable percentage of human but not mouse OPC. For myelinating oligodendrocytes (MOL) see **Figure 3B**.

**Supplemental Figure S4.**
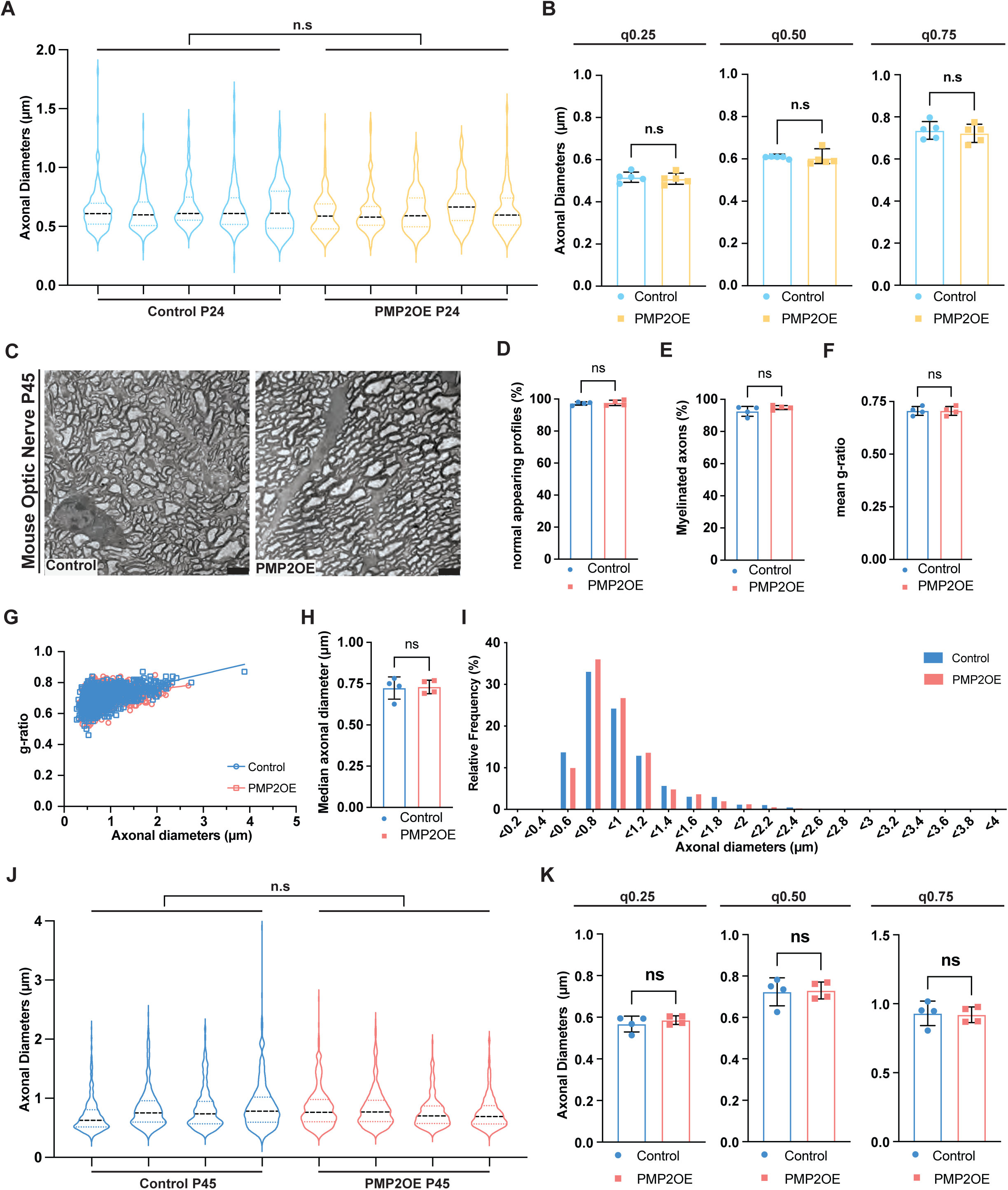
Analysis of diameters of CNS axons in PMPOE and control mice. **A** Violin plot showing diameters of axons in the optic nerves of control versus PMP2OE mice at age P24 as assessed on images as in **Figure 5F**. Each violin represents one out of five control mice and five PMP2OE mice at age P24; Horizontal lines indicate the 25th, 50th (median), and 75th percentiles to show spread and skewness. For median axonal diameters see **Figure 5K** and for frequency distribution see **Figure 5L**. Evaluation of the distribution of the diameters using quantile analysis (lqmm) did not identify differences at the quartile boundaries (first and second quartile boundary: p = 0.06274; median: p = 0.2071; third and fourth quartile boundary: p = 0.2071). **B** Bar graphs showing comparison of means across quartile groups (q25, 25%; q50, 50%; q75, 75%) of axon diameters in the optic nerves of each mouse for control (blue) or PMP2OE (yellow) at age P24 as assessed on images as in **Figure 5F**. Each datapoint represents one individual. Bar graph gives mean ±SD; q25 n.s. p=0.6586, q50 n.s. p=0.8005, q75 n.s. p=0.6282 by Two-tailed Student’s t-test. Note that the diameters of axons do not differ between PMP2OE and control mice. **C-K** Electron micrographs of cross-sectioned optic nerves dissected from control and PMP2OE mice at P45 **(C)** and genotype-dependent quantification **(D-K)**. **C** Images representative of n=4 mice per genotype. Scale bar 2.5 μm. **D** Quantification of non-pathological axon/myelin-units on images as in **C** reveals no significant difference between control and PMP2OE mice at P45 (n=4). Mean ±SD; datapoints represent individual mice; n.s. p>0.05 by Two-tailed Student’s t-test. **E** Quantification of the percentage of axons that are myelinated on images as in **C** reveals no significant difference between control and PMP2OE mice at P45 (n=4). Mean ±SD; datapoints represent individual mice; n.s. p>0.05 by Two-tailed Student’s t-test. **F,G** g-ratio analysis of normal appearing axon-myelin profiles in optic nerves reveals a normal g-ratio suggesting appropriately thick myelin sheaths in PMP2OE compared to Ctrl mice at P45 (n=4). **F** Bar graph gives mean ±SD; n.s. p>0.05 by Two-tailed Student’s t-test. **G** Scatter plot displays g-ratio and respective axonal diameters; datapoints show 399-481 axon/myelin-units per mouse. **H,I** Genotype-dependent quantification reveals similar axonal diameters in optic nerves of PMP2OE and Ctrl mice at P45 (n=4). **H** Bar graph gives median ±SD; n.s. p>0.05 by Two-tailed Student’s t-test. **I** No frequency distribution shift in PMP2OE mice compared to controls. Data presented as frequency distribution with 0.2 μm bin width; 1782 axons in n = 4 control and 1796 axons in n = 4 PMP2OE mice. **J** Violin plot showing diameters of axons in the optic nerves of control (blue) or PMP2OE (red) mice at age P45 as assessed on images as in **supplemental Figure S4C**. Each violin represents one mouse at age P45; Horizontal lines indicate the 25th, 50th (median), and 75th quantiles to show spread and skewness. Evaluation of the distribution of the diameters using quantile analysis (lqmm) did not identify differences at the quartile boundaries (first and second quartile boundary: p = 0.2932; median: p = 0.9293; third and fourth quartile boundary: p = 0.628). **K** Bar graphs showing comparison of means across quartile groups (q25, 25%; q50, 50%; q75, 75%) of axon diameters in the optic nerves of each mouse for control (blue) or PMP2OE (red) mice at age P45 as assessed on images as in **supplemental Figure S4C.** Each datapoint represents one individual. Bar graph gives mean ±SD; q25 n.s. p=0.4205, q50 n.s. p=0.8694, q75 n.s. p=0.8558 by Two-tailed Student’s t-test. Note that the diameters of axons do not differ between PMP2OE and control mice.

**Supplemental Figure S5.**
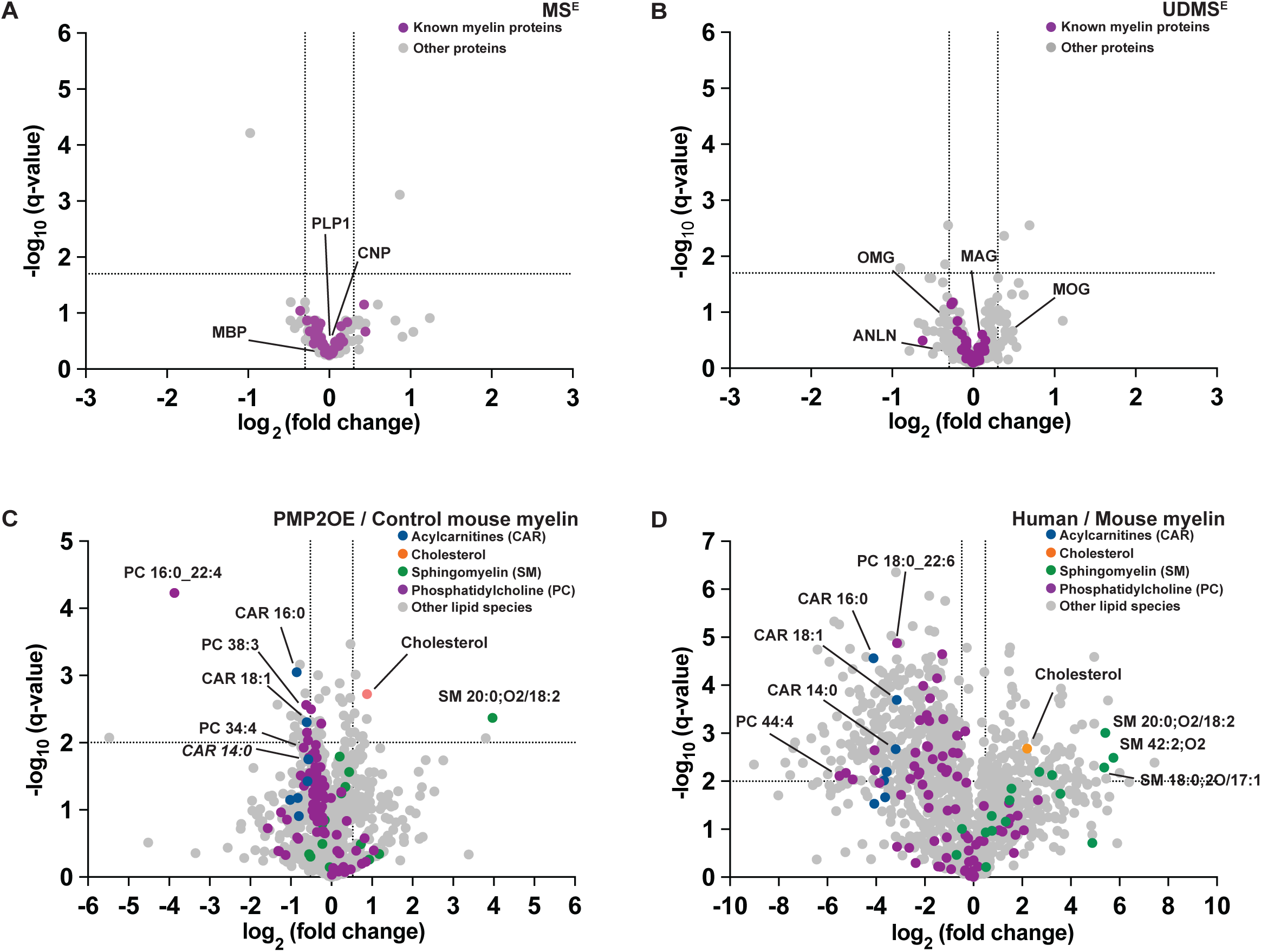
Volcano plots summarizing myelin proteome and lipidome analysis. **A,B** Volcano plots summarizing genotype-dependent comparison as measured by two different mass spectrometric acquisition modes (MS^E^ in **A,** UDMS^E^ in **B**) comparing myelin purified from the brains of PMP2OE and control mice at age P45. Datapoints represent relative abundance of proteins quantified in myelin of PMP2OE compared to controls and are plotted as the log_2_-transformed fold-change (FC) on the x-axis against the −log_10_-transformed q-value on the y-axis. Vertical stippled lines mark a log_2_-fold change of 0.5 or −0.5 threshold of changed protein abundance; horizontal stippled lines indicate a −log_10_-transformed q-value of 2 reflecting a q-value of 0.01 as significance threshold. Datapoints highlighted in magenta represent known myelin proteins; all other datapoints (gray) reflect other proteins in the dataset. In **A,** datapoints highlighted in magenta indicate exceptionally abundant myelin proteins PLP, MBP, and CNP; other datapoints represent other proteins identified in myelin. In **B**, datapoints highlighted in magenta indicate known myelin marker proteins MOG, MAG, OMG, and ANLN; other datapoints represent other proteins identified in myelin. Note that no known myelin protein displays significantly different abundance between genotypes. PMP2 is not displayed as it was not detected in Ctrl myelin. For heatmap highlighting known myelin proteins see **Figure 6A**. For entire dataset see **Supplemental Data Table S1**. **C** Volcano plot summarizing genotype-dependent comparison of lipid species as measured by mass spectrometry in positive ionization mode comparing myelin purified from the brains of PMP2OE and control mice at age P45. Datapoints represent relative abundance of lipid species quantified in myelin of PMP2OE compared to controls and are plotted as the log_2_-transformed fold-change (FC) on the x-axis against the −log_10_-transformed q-value on the y-axis. Vertical stippled lines mark a log_2_-fold change of 0.5 or −0.5 threshold of changed lipid species abundance; horizontal stippled lines indicate a −log_10_-transformed q-value of 2 reflecting a q-value of 0.01 as significance threshold. Indicated selected lipid species are highlighted as colored datapoints; all other datapoints (gray) reflect other lipid species in the dataset. For lipid classes see **Figure 6B**. For dataset see **Supplemental Data Table S2**. **D** Volcano plot summarizing genotype-dependent comparison of lipid species as measured by mass spectrometry in positive ionization mode comparing myelin purified from the brains of human individuals and mice. Datapoints represent relative abundance of lipid species quantified in myelin of humans compared to mice and are plotted as the log_2_-transformed fold-change (FC) on the x-axis against the −log_10_-transformed q-value on the y-axis. Vertical stippled lines mark a log_2_-fold change of 0.5 or −0.5 threshold of changed lipid species abundance; horizontal stippled lines indicate a −log_10_-transformed q-value of 2 reflecting a q-value of 0.01 as significance threshold. Indicated selected lipid species are highlighted as colored datapoints; all other datapoints (gray) reflect other lipid species in the dataset. For lipid classes see **Figure 6C**. For dataset see **Supplemental Data Table S2.**

